# Predicting the topography of fitness landscapes from the structure of genotype-phenotype maps

**DOI:** 10.1101/2025.02.14.638275

**Authors:** Malvika Srivastava, Ard A. Louis, Nora S. Martin

## Abstract

Ruggedness, the prevalence of fitness peaks, and navigability, the existence of fitness-increasing paths to a target, are key factors affecting evolution on fitness landscapes. We analytically predict landscape ruggedness for genotype-phenotype maps with randomly assigned fitness, using only the sizes of neutral components – mutationally connected genotype sets sharing the same phenotype– and their evolvabilities, the number of neighbouring phenotypes. Further, more evolvable peaks tend to have higher fitness and a minimal evolvability is required for navigability.

---

Fitness landscapes assign fitness values to genotypes. Their topography is central for predicting evolutionary outcomes since this sets the fitness-increasing, ‘accessible’ paths, along which populations, especially those described by the strong-selection-weak-mutation regime, traverse the landscape [1]. Important topographical features include the prevalence of local optima or *peaks* (here referred to as *ruggedness*, but the term is used for several related concepts [2]). In addition *navigability* quantifies the probability that an accessible path to the global optimum exists [3].

Fitness landscapes are exponentially large, typically with *K*^*L*^ genotypes for a sequence length L and alphabet size *K*. Since partial analyses may misidentify saddle points as peaks [6], exhaustive large-scale analyses are essential. This can be achieved with computational genotype to fitness models such as the House-of-Cards (HoC) model, which randomly allocates fitness values to genotypes and permits analytic calculations [1, 7]. While these models capture the high dimensionality of fitness landscapes [1], they abstract away other biological details.

A more biologically motivated approach is to start with a computational or empirical genotype-phenotype (GP) map, and then to assign a fitness value to each phenotype [3], thus creating a genotype-phenotype-fitness (GPF) map. Such a landscape inherits the properties of the GP map, many of which are thought to be ‘universal’ [8, 9].

One biological property captured by GP maps is the prevalence of neutral mutations [10], which preserve phenotype and fitness, thus enabling accessible paths [11, 12]. Neutral mutations can connect genotypes mapping to the same phenotype to form large *neutral components* (NCs) [4, 10]. Larger NCs tend to have higher *evolvability*, measured by the number of unique phenotypes among the mutational neighbours of the NC [4, 13] (note that the term ‘evolvability’ is also used for other quantities [14]). Numerical analyses have suggested that these and further universal properties result in topographical features such as low ruggedness and high navigability in many GPF maps, even when fitness values are randomly assigned to phenotypes [3], but the exact causal and quantitative links were previously unclear.

In this paper, we derive and test analytic predictions for the topography of GPF maps with a random phenotype-fitness (PF) map (Fig 1A). We show that the prevalence of fitness peaks is completely determined by NC sizes and evolvabilities. Secondly, we find that the higher the evolvability, the higher the expected fitness of a peak. Finally, we estimate the number of accessible paths, finding that low-evolvability GP maps are unlikely to be navigable.

**FIG. 1.**
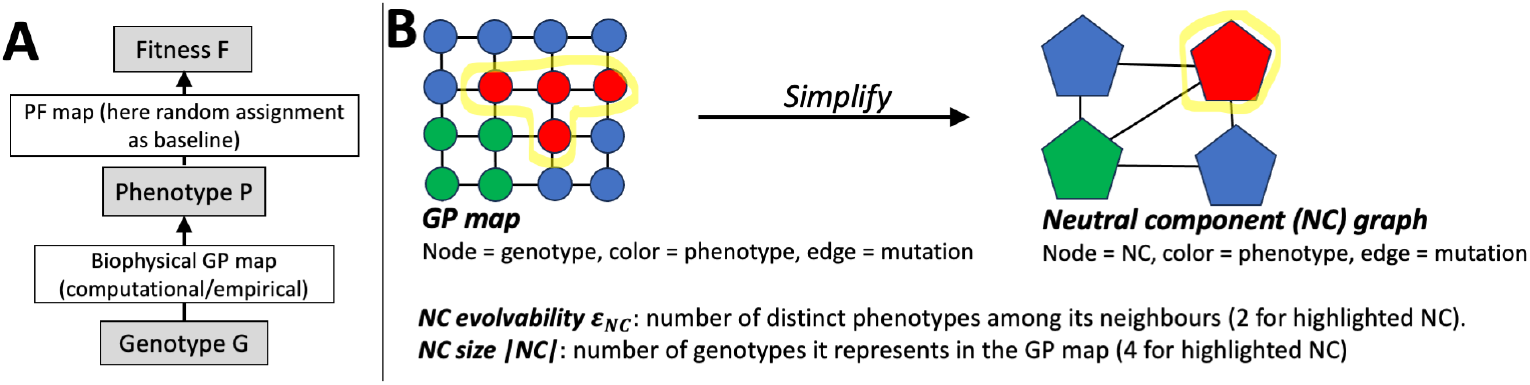
GPF maps and the NC graph: (A) In a GPF map, each genotype is first mapped to a phenotype using a biophysical GP map, and then to fitness using a PF map (in this case, a random assignment) [3]. (B) Each NC represents a set of mutationally connected genotypes that share the same phenotype [4] and thus have a single associated fitness value. The study of ruggedness and navigability in GPF maps can be simplified by reducing the underlying GP map (left) to a NC graph (right) [5], which is typically orders of magnitude smaller.

We validate our predictions using simulations on a diverse set of nine GP maps. These include six biophysical computational GP maps from ref [3], describing RNA secondary structure (“RNA structure”), protein quaternary structure self-assembly (“self-assembly”), and hydrophobic-polar protein folding (“HP x,” with x indicating system size). Additionally, we build a GP map, where each genotype maps to the set of specifically binding RNA-binding proteins (note that this is a highly simplified way of defining a simple, categorical phenotype per genotype), based on empirical binding data curated in ref [15]. Lastly, we employ an abstract GP map model (“Fibonacci”) and its low-evolvability counterpart (“LE Fibonacci”), both from ref [16].

## Landscape topography follows from a coarse-grained NC graph representation of the GP map

To make the analysis tractable, we work with GPF maps at the coarse-grained level of neutral components (NCs) (similar to existing coarse-grained treatments [5, 17, 18]). Mutational paths within a NC are accessible by definition, and so we can sidestep the complexity of exhaustively analysing these paths by abstracting each NC to a single entity. This means rewriting the GP map as a network of NCs, the *NC graph*: Each NC is a node, and two nodes are connected by an edge if there is a mutational connection between the two NCs in the GP map (Fig 1B) [5]. Even though such a NC graph is orders of magnitude smaller than the full GP map in our examples, it contains all information needed to identify accessible paths and peaks: A NC is a peak if all its mutational neighbours on the *NC graph* are lower in fitness. Similarly, an accessible path from an initial NC to a final NC exists on the GPF map, if and only if such a path exists in the NC graph [5].

## Ruggedness is set by the number, size and evolvability of NCs

For a NC to be a peak, it must be fitter than its *ϵ*_*NC*_ distinct phenotypic neighbours. Upon random fitness assignment to phenotypes, any fitness ranking between the NC’s phenotype and its *ϵ*_*NC*_ neighbours is equally likely, and thus the likelihood that a NC is a peak decreases rapidly with its evolvability as 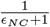 (as in HoC model calculations [7], but at the coarse-grained level of NC graph nodes rather than individual genotypes.). The expected ruggedness, defined here as the fraction of all *genotypes* in a GPF map that are in peaks, can be calculated as:

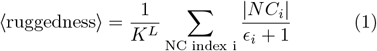

where |*NC*_*i*_| is the number of genotypes in *NC*_*i*_, and we normalise by the total number of genotypes *K*^*L*^.Thus, for a random phenotype-fitness assignment without ties, the only factors determining ruggedness are the set of NC sizes {|*NC*_*i*_|} and NC evolvabilities {*ϵ*_*i*_}. Other properties of GP maps (for example, dimensionality [3], see SM section S2.2.1) only affect the number of peaks indirectly insofar as they influence the NCs and their evolvabilities.

Fig 2 demonstrates that eq. (1) accurately predicts the mean ruggedness found in simulations for all nine GP map examples. Ruggedness is low across the GP maps and lower than in a randomised ‘HoC-like’ landscape of the same size (see SM section S2.2.1), with two exceptions: the LE Fibonacci model, which was specifically designed as a low-evolvability test case, and the self-assembly model, where the short sequence length chosen for computational reasons leads to a particularly low number of phenotypes [3] and thus limits evolvability. Eq. (1) allows us to see how this low ruggedness follows from the structure of the GP map: higher NC evolvabilities imply lower ruggedness, and for a given set of evolvability values, ruggedness will be lowest if the largest NCs have highest evolvabilities, i.e. if there is a a positive correlation between NC size and evolvability (see for example our GP maps in SM figure S4).

**FIG. 2.**
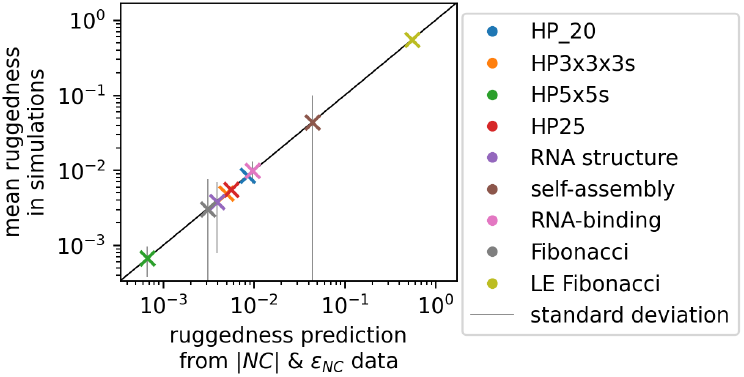
Ruggedness, the fraction of genotypes in peaks, can be predicted from the set of NC sizes and evolvabilities: For our nine GP maps, the mean ruggedness in simulations (y-axis, average over 10^3^ PF maps, error bars show standard deviation) is well-predicted by eq. (1) (x-axis).

## Evolvability-enhancing epistasis can reduce ruggedness

Since a positive correlation between NC size and evolvability is one key factor for low ruggedness, we turn to the roots of this correlation. Although all genotypes in a NC generate the same phenotype, high evolvability requires that these genotypes have many different alternative phenotypes in their mutational neighbourhoods [16]. Shifts in mutational effects due to genetic background are known as epistasis, and we will call this specific form *evolvability-enhancing epistasis* ^1^. Its significance is illustrated by the LE Fibonacci map, which lacks this epistasis by construction [16] and has a much higher ruggedness than the standard Fibonacci map (Fig 2).

## Higher-evolvability peaks tend to be higher in fitness

High-evolvability peaks tend to have high fitness, since these NCs are only peaks if they outcompete many mutational neighbours. For example, if the PF map is drawn from a uniform distribution between 0 and 1, the mean peak fitness ⟨*F*⟩ _peak_ and its variance Var(*F*_peak_) depend on the peak’s evolvability *ϵ*_*NC*_ as (see SM section S1.2):

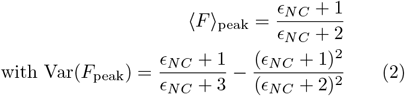

These predictions agree with computational results (Fig. 3A), and a similar trend holds for PF maps drawn from an exponential distribution, where fitness has no fixed upper limit (Fig. 3B, analytic expression in section S1.2 of the SM). Since evolvability usually increases with NC size [4, 13], this implies that large peaks, which are more likely to arise in evolutionary processes [19], are likely high in fitness (section S2.3 in the SM).

**FIG. 3.**
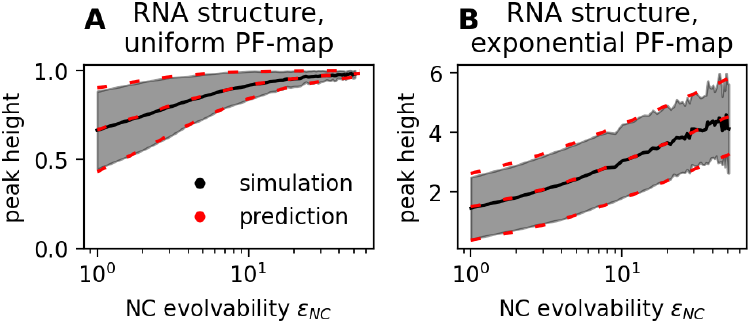
More evolvable peaks have higher fitness: For the simulated RNA GPF maps, peak fitness is plotted against evolvability (mean and standard deviation in grey) for 10^3^ PF map realisations. High-evolvability peaks are higher in fitness, in agreement with our analytic predictions (mean and standard deviation as dashed red lines). (A) PF map drawn from a uniform distribution (see eq. (2)). (B) PF map drawn from an exponential distribution (see eqs (6&8) in the SM).

## Minimum evolvability required for navigability in simple GP maps

Given that GPF maps are multi-peaked, a key question is whether they nevertheless have fitness-increasing paths to a target. This is quantified by the *navigability*, the likelihood that at least one accessible path exists for a randomly chosen source-target pair when the target fitness is set to the maximum fitness value [3].

While navigability is harder to predict from simple NC characteristics than ruggedness, we can make progress for a simple case: GP maps with a single NC per phenotype, where each NC connects to any of the *n*_*p*_ phenotypes with a constant probability 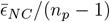. In this scenario, a NC graph is expected to be non-navigable if (note the caveats in the derivation in SM section S1.3):

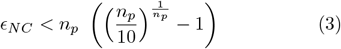

While GP maps exceeding this evolvability are not guaranteed to be navigable, eq. (3) predicts that a highly connected, high-evolvability NC graph is a minimum requirement for the existence of accessible paths, as hypothesised in ref [5].

To apply eq (3), which assumes a single NC per phenotype, to our GP maps with multiple NCs per phenotype, we computed evolvabilities in two simplified versions of each GP map: (1) connecting all NCs of a phenotype through mutations or (2) retaining only the largest NC per phenotype. These simplified maps are used to compute evolvabilities, but the navigability simulations use the full map. Fig 4 shows that even with these simplifications and despite relying on strong approximations, eq 3 successfully identifies a low-evolvability regime in which GP maps are non-navigable. Further applications of eq 3, including to GP maps with different evolvability distributions, are given in the SM section S2.4.

**FIG. 4.**
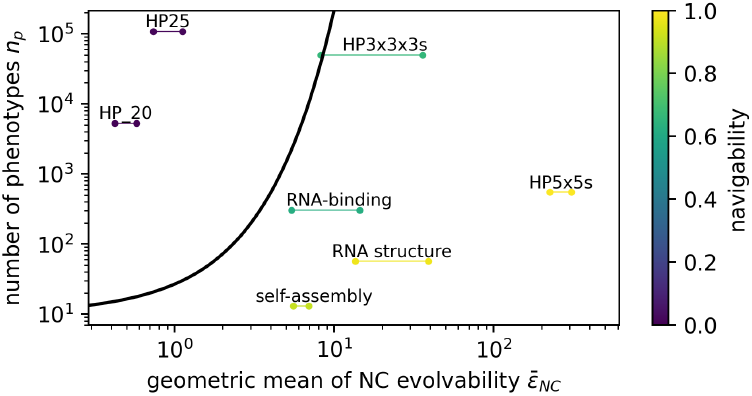
Evolvability plays a key role for navigability: For each GP map, the number of phenotypes is plotted against the geometric mean of its evolvabilities (since a single *ϵ*_*NC*_ = 0 would bring the geometric mean to zero, this is treated as *ϵ*_*NC*_ = 0.01), with navigability shown on the colour scale. Each map is represented by two evolvability values, corresponding to two ways of resolving NC fragmentation. Eq. (3) (black line) successfully identifies an evolvability below which GP maps are non-navigable, despite relying on strong approximations. The Fibonacci model is excluded and analysed in SM section S2.4.2.

## Conclusions

To sum up, we analyse ruggedness and navigability in GPF maps with a simplified NC graph [5], which abstracts away the sequence space of the full GP map, while still accounting for indirect paths, dimensionality and neutral mutations. For random phenotype-fitness maps, this NC graph representation enabled us to make analytic predictions. Ruggedness only depends on the NC sizes and evolvabilities, with other GP map features affecting ruggedness only indirectly through their influence on these two NC properties. Further, the prevalence of peaks in GPF maps decreases when NC size and evolvability are positively correlated, which holds across well-studied GP maps (at least for *phenotypic* evolvability [8, 9, 13], and likely also for NCs [4]) and is due to a special type of evolvability-enhancing epistasis [16]. However, other types of epistasis can fragment NCs [4], increasing ruggedness. The interplay between epistasis and ruggedness in GP maps remains intricate and context-dependent.

Secondly, we found that high-evolvability NCs, if they form peaks, have exceptionally high fitness. These high-evolvability NCs are known to be large [4, 13] and thus more likely to arise as potential variation in evolutionary processes [19].

Finally, we turned to accessible paths and derived a scaling that identifies high evolvability as a minimum requirement for navigability, and, despite relying on strong approximations, successfully identifies non-navigable GP maps in our dataset.

Many of our GPF maps, despite their rugged ‘House-of-Cards’-like PF map, have high navigability, and their large NCs have high evolvabilities, making them unlikely to be low-fitness peaks. Thus, such GPF maps may add a qualitatively new contribution to a broader discussion of “rugged yet […] navigable” landscapes [20–23].

While some GPF map calculations may superficially resemble calculations for HoC GF maps (for example, eq. (1) would equal the HoC expression if we replaced *ϵ* by (*K* − 1)*L* [7] and peaks are higher for larger *L* [24]), there is a key difference: In contrast to GF maps, where the mutational neighbourhood size (*K* − 1)*L* is usually the same for all genotypes, in GPF maps, the relevant mutational neighbourhood size is the evolvability *ϵ*, which can vary by orders of magnitude *within* a single GP map (see SM Fig. S4). This leads to a richer phenomenology, with NCs having vastly different sizes and evolvabilities, and different probabilities of being peaks and expected fitness if they are peaks.

An interesting extension would go beyond the random PF map and consider more realistic, correlated PF maps where similar phenotypes have similar fitness values [3]. Future research should leverage the computational and conceptual simplifications of the NC graph to study such PF maps, as well as the likelihood of an accessible path being realised in an evolutionary process. This could involve incorporating mutation probabilities linking different NCs [19, 25] into the NC graph as edge weights (see weighted graphs in [17, 18]). Finally, the effectiveness of using the NC graph for this analysis suggests it could help provide useful quantitative insights into other aspects of GP map structure and their evolutionary consequences.

## METHODS

### NC graph construction from GP maps

We identified NCs using a decomposition algorithm [26], and iterated through all mutations on all genotypes to obtain all mutational connections between NCs. With this information, we constructed the NC graph, recording the evolvability *ϵ*_*NC*_ and size |*NC*| of each NC. NCs with phenotype ‘deleterious’ (for example, non-terminating assemblies) are assumed to have zero fitness [3], such that they play no role in accessible paths and are thus not included in the NC graph.

### Ruggedness simulations

For each realisation of the random map, we draw one fitness value per phenotype from a uniform distribution from 0 to 1 (except Fig. 3B - exponential distribution with *λ* = 1). To find the number of genotypes in peaks, we sum the sizes |*NC*| of all NCs whose fitness exceeds that of all neighbouring NCs. Ruggedness data is based on 10^3^ PF map realisations.

### Navigability simulations

We pick 10 phenotypic source-target pairs each for 100 PF maps drawn from a uniform distribution, with the target’s fitness set to 1. We chose a starting NC at random from all NCs mapping to the source phenotype, with each NC being picked with a probability proportional to its size |*NC*| (to match the navigability definition [3], where the start is chosen uniformly from all genotypes mapping to the source phenotype). Then, we look for accessible paths leading to any NC of the target phenotype, using standard graph algorithms [27] on a directed NC graph, where edges point in fitness-increasing direction (similar to the component network algorithm for navigability in ref [5]).

### GP map data

For the HP, RNA and self-assembly data, we use the GP maps from Greenbury et al. [3] (the datasets labelled ‘HP *’, ‘RNA 12’ and ‘S 2 8’ respectively).

The RNA-binding map is based on a curated empirical dataset [15] of RNAcompete data for *Drosophila Melanogaster*. To obtain a categorical, many-to-one GP map, we map each length-seven RNA genotype to the *set of* specifically binding RNA binding proteins (RBPs) as a phenotype (i.e. those with an enrichment score of *>* 0.35 as in [15]), and treated sequences without any RBP as deleterious. Note that this GP map would change if data for more RBPs were added. Further, including all RBP without focusing on specific cell types and neglecting binding strengths is a highly simplified treatment.

The Fibonacci GP map we use is Weiß and Ahnert’s [16] generalisation of the original [28]: genotypes are length-*L* strings made up of *K* distinct letters, with one letter acting as a stop codon (here *L* = 12, *K* = 3). The phenotype is obtained by cutting the genotype at the first stop codon, and genotypes without stop codons are undefined/deleterious. In the low-evolvability version, the first stop codon cannot mutate [16]. To ensure symmetry of allowed mutations, we additionally exclude mutations that introduce a new stop codon before the first stop codon (if ‘B’ is the stop codon, ‘ABAB’ cannot mutate to ‘AAAB’, so the reverse mutation ‘AAAB’ to ‘ABAB’ is also forbidden).

## DATA AVAILABILITY

Our code is available at https://github.com/noramartin/gpf_maps/.

## ACKNOWLEDGMENTS

This research was funded by the Swiss National Science Foundation (Grant numbers PP00P3 170604 and the PhD mobility grant), the Spanish Ministry of Science and Innovation through the Centro de Excelencia Severo Ochoa (CEX2020-001049-S, MCIN/AEI/10.13039/501100011033), the EMBL partnership, the Generalitat de Catalunya through the CERCA programme, and Grant JDC2022-049526-I funded by MCIN/AEI/10.13039/501100011033 and by European Union NextGenerationEU/PRTR. We thank J. L. Payne for providing the RBP data and S.F. Greenbury for the computational GP map data.

## Footnotes

1 While the original paper preferred ‘non-local mutation effects’ [16], here we will use the more general term ‘epistasis’.

