## Supplementary text for "Predicting the topography of fitness landscapes from the structure of genotype-phenotype maps"

#### S1 Analytic derivations

##### S1.1 Ruggedness derivation - additional information

In our ruggedness calculation, we used the following argument: a NC has to be the fittest among its $\epsilon_{NC}$ phenotypic neighbours to be a peak. Thus, in a random PF map, a NC with evolvability $\epsilon_{NC}$ is a peak with probability (similar to a HoC model [1], but with neighbourhood size $\epsilon_{NC}$):

$$P(\text{is peak}) = 1/(\epsilon_{NC} + 1) \tag{1}$$

Thus, we only need to sum over all NCs and multiply by the number of genotypes in each NC ($|NC|$) to get the expected prevalence of peaks:

$$\langle \text{fraction genotypes in peaks} \rangle = \frac{1}{K^L} \sum_{\text{NC indices } i} \frac{|NC_i|}{\epsilon_i + 1} \tag{2}$$

Here, $K$ is the alphabet size and $L$ the sequence length, such that the normalisation $N_G = K^L$ is the total number of genotypes. Ruggedness will decrease if the mutational connectivity of the NC graph is raised by increasing $\epsilon_i$, which is intuitive and in line with previous work on specific classes of evolutionary networks [2].

The following calculation shows that this eq. (2) is valid despite the fact that individual summands in eq. (2) are not independent: if the $i^{th}$ NC is a peak, then none of its $\epsilon_i$ mutational neighbours can be peaks. These interdependencies can be ignored when estimating the *average* ruggedness, where the average is computed over many PF maps:

$$\begin{aligned}
\langle \text{fraction genotypes in peaks} \rangle &= \frac{1}{K^L} \langle \sum_{\text{NC indices } i} |NC_i| \text{ if } NC_i \text{ is peak under this PF map, else } 0 \rangle \\
&= \frac{1}{K^L} \sum_{\text{NC indices } i} \langle |NC_i| \text{ if } NC_i \text{ is peak under this PF map, else } 0 \rangle \\
&= \frac{1}{K^L} \sum_{\text{NC indices } i} |NC_i| \times \langle 1 \text{ if } NC_i \text{ is peak under this PF map, else } 0 \rangle \\
&= \frac{1}{K^L} \sum_{\text{NC indices } i} |NC_i| \times \text{fraction of PF maps for which } NC_i \text{ is a peak} \\
&= \frac{1}{K^L} \sum_{\text{NC indices } i} \frac{|NC_i|}{\epsilon_i + 1}
\end{aligned}$$

Thus, we can compute the average as in eq. (2) without considering dependencies between different NCs in a specific realisation of the PF map.

##### S1.2 Derivation: expected height of a peak with evolvability $\epsilon_{NC}$

Next, we turn to the expected fitness of a peak of evolvability $\epsilon_{NC}$. Using Bayes' rule, we can write the probability that a NC of evolvability $\epsilon_{NC}$ has fitness $F$, given that it is a peak, as:

$$P(F|\text{is peak}) = P_F(F) \frac{P(\text{is peak}|F)}{P(\text{is peak})} \tag{3}$$

We know that $P(\text{is peak}) = (1+\epsilon_{NC})^{-1}$, regardless of the PF map (see eq. (1)). $P_F(F)$ and $P(\text{is peak}|F)$ will depend on the distribution underlying the PF map, $P_F(F)$.

Alternatively, we can reduce the problem to standard calculations on order statistics: Whenever a NC of evolvability $\epsilon_{NC}$ is a peak, it is the highest of $\epsilon_{NC}+1$ relevant phenotypes (itself and its $\epsilon_{NC}$ mutational neighbours). Thus, its fitness follows the same distribution as the maximum value obtained after $\epsilon_{NC}+1$ draws from $P_F(F)$ (i.e. the highest order statistic after $n = \epsilon_{NC}+1$ draws). This connection to order statistics means that we could simply look up eq. (3) [3], as well as the mean and standard deviation of $F$, for uniform [3] and exponential [4] PF maps. Nevertheless, we will give full derivations for completeness in the following sections.

##### S1.2.1 PF map drawn from a uniform distribution

If the PF map is sampled from a uniform distribution between 0 and 1, we have $P_F(F) = 1$ for $0 < F < 1$. The probability $P(\text{is peak}|F)$ is the likelihood that all $\epsilon_{NC}$ phenotypes in the neighbourhood have fitness values less than $F$, which for a uniform distribution is $F^{\epsilon_{NC}}$. Thus, we have:

$$P(F|\text{is peak}) = (1+\epsilon_{NC})F^{\epsilon_{NC}} \quad \text{for } 0 < F < 1$$

To obtain the mean of this height distribution, we write:

$$\begin{aligned}\langle F\rangle_{\text{peak}} &= \int_0^1 dF \cdot F \cdot P(F|\text{is peak}) \\ &= \int_0^1 dF (1+\epsilon_{NC}) \cdot F^{\epsilon_{NC}+1} \\ &= \frac{\epsilon_{NC}+1}{\epsilon_{NC}+2} \end{aligned} \tag{4}$$

To obtain the variance of this height distribution, we write:

$$\begin{aligned}\text{Var}(F_{\text{peak}}) &= \langle F^2\rangle_{\text{peak}} - \langle F\rangle^2_{\text{peak}} \\ &= \int_0^1 dF \cdot F^2 \cdot P(F|\text{is peak}) - \langle F\rangle^2_{\text{peak}} \\ &= \int_0^1 dF\ (1+\epsilon_{NC})\ F^{(\epsilon_{NC}+2)} - \langle F\rangle^2_{\text{peak}} \\ &= \frac{\epsilon_{NC}+1}{\epsilon_{NC}+3} - \left(\frac{\epsilon_{NC}+1}{\epsilon_{NC}+2}\right)^2 \end{aligned} \tag{5}$$

As expected, these expressions for the mean and variance agree with the literature on order statistics of uniform distributions: they are the mean and the variance of the maximum after $\epsilon_{NC}+1$ draws [3].

##### S1.2.2 PF map drawn from an exponential distribution

For a PF map sampled from an exponential distribution, the fitness distribution is given by $P_F(F) = \lambda \exp(-\lambda F)$. Then, $P(\text{is peak}|F) = (1-\exp(-\lambda F))^{\epsilon_{NC}}$ because the probability that all $\epsilon_{NC}$ neighbours have a lower fitness than the peak NC is $[P(F_i < F)]^{\epsilon_{NC}} = [\int_0^F dx \cdot \lambda \exp(-\lambda x)]^{\epsilon_{NC}} = (1-\exp(-\lambda F))^{\epsilon_{NC}}$.

Thus, we have:

$$P(F|\text{is peak}) = (1+\epsilon_{NC})\lambda \exp(-\lambda F)\cdot(1-\exp(-\lambda F))^{\epsilon_{NC}}$$

To obtain the mean of this height distribution, we write:

$$\begin{aligned}\langle F\rangle_{\text{peak}} &= \int_0^\infty dF\cdot F\cdot P(F|\text{is peak})\\ &= (1+\epsilon_{NC})\cdot\lambda\int_0^\infty dF\cdot F\cdot\exp(-\lambda F)\cdot(1-\exp(-\lambda F))^{\epsilon_{NC}}\end{aligned}$$

Substituting $e^{-\lambda F} = t \implies dF = -dt/(\lambda e^{-\lambda F}) = -dt/(\lambda t)$ and $F = -\ln(t)/\lambda$, we get:

$$\begin{aligned}\langle F\rangle_{peak} &= \frac{(\epsilon_{NC}+1)}{\lambda}\int_1^0 (1-t)^{\epsilon_{NC}}\cdot\ln(t)\cdot dt\\ &= -\frac{(\epsilon_{NC}+1)}{\lambda}\int_0^1 (1-t)^{\epsilon_{NC}}\cdot\ln(t)\cdot dt\end{aligned}$$

Upon integrating by parts, we get:

$$\begin{aligned}\langle F\rangle_{peak} &= \frac{(\epsilon_{NC}+1)}{\lambda}\left[\ln(t)\cdot\frac{(1-t)^{(\epsilon_{NC}+1)}}{(\epsilon_{NC}+1)}\Bigg|_0^1 - \int_0^1 dt\frac{1}{t}\frac{(1-t)^{(\epsilon_{NC}+1)}}{(\epsilon_{NC}+1)}\right]\\ &= \frac{1}{\lambda}\left[\ln(t)\cdot(1-t)^{(\epsilon_{NC}+1)}\Big|_0^1 + \int_0^1 dt\frac{1-(1-t)^{(\epsilon_{NC}+1)}}{1-(1-t)} - \int_0^1 dt\frac{1}{t}\right]\end{aligned}$$

The first and the third term of the above equation give rise to $\ln(0)$ terms that cancel. Then, we are left with one term, which can be simplified by substituting $s = 1-t$:

$$\begin{aligned}\langle F\rangle_{peak} &= \frac{1}{\lambda}\int_1^0 -ds\frac{1-s^{(\epsilon_{NC}+1)}}{1-s}\\ &= \frac{1}{\lambda}\int_0^1 ds\frac{1-s^{(\epsilon_{NC}+1)}}{1-s}\end{aligned}$$

Now, using the fact that the Harmonic number $H_n := \int_0^1 dx\frac{1-x^n}{1-x}$, we have:

$$\langle F\rangle_{peak} = \frac{H_{\epsilon_{NC}+1}}{\lambda} \tag{6}$$

Similarly, to obtain the variance of this height distribution, we write:

$$\text{Var}(F_{\text{peak}}) = \langle F^2\rangle_{\text{peak}} - \langle F\rangle^2_{\text{peak}}$$

To calculate $\langle F^2\rangle_{\text{peak}}$, we will rely on cumulative distribution functions to simplify the calculations. To obtain the cumulative distribution, we simply integrate $P(F|\text{is peak})$ to get $P(F < f|\text{is peak}) = \int_0^f(1+\epsilon_{NC})\lambda\exp(-\lambda F)\cdot(1-\exp(-\lambda F))^{\epsilon_{NC}} = (1-\exp(-\lambda f))^{\epsilon_{NC}+1}$. We can compute $\langle F^2\rangle_{\text{peak}}$ from this CDF using $\langle X^2\rangle = \int_0^\infty dx\cdot 2x(1-F_X(x))$ (ref [5] with $F \geq 0$):

$$\langle F^2 \rangle_{\text{peak}} = \int_0^\infty dF \cdot 2F \cdot (1 - (1 - \exp(-\lambda F))^{\epsilon_{NC}+1})$$

Substituting $\lambda F = -\log(1-u)$ gives:

$$\langle F^2 \rangle_{\text{peak}} = \frac{-2}{\lambda^2} \int_0^1 du \cdot \log(1-u) \cdot \frac{1 - u^{\epsilon_{NC}+1}}{1-u}$$

Now using the fact that $\frac{1-u^n}{1-u} = \sum_{k=0}^{n-1} u^k$, expanding $\log(1-u) = -\sum_{m=1}^{\infty} \frac{u^m}{m}$ gives:

$$\langle F^2 \rangle_{\text{peak}} = \frac{2}{\lambda^2} \cdot \sum_{m=1}^{\infty} \sum_{k=0}^{\epsilon_{NC}} \int_0^1 du \frac{u^{m+k}}{m}$$

Now integrating over $u$ gives:

$$\begin{aligned}
\langle F^2 \rangle_{\text{peak}} &= \frac{2}{\lambda^2} \cdot \sum_{m=1}^{\infty} \sum_{k=}^{\epsilon_{NC}} \frac{1}{m(m+k+1)} \\
&= \frac{2}{\lambda^2} \cdot \sum_{m=1}^{\infty} \sum_{k'=1}^{\epsilon_{NC}+1} \frac{1}{m(m+k')}
\end{aligned}$$

Where the dummy index on the sum was changed to $k' = k+1$.

Finally, re-arranging the terms and using the definition of the Harmonic number $H_n = \sum_{k=1}^{n} \frac{1}{k}$ gives:

$$\begin{aligned}
\text{Var}(F_{\text{peak}}) &= \frac{1}{\lambda^2} \cdot \sum_{k=1}^{\epsilon_{NC}+1} \frac{2}{k} \sum_{m=1}^{\infty} \left( \frac{1}{m} - \frac{1}{m+k} \right) - \langle F \rangle_{\text{peak}}^2 \\
&= \frac{1}{\lambda^2} \cdot \sum_{k=1}^{\epsilon_{NC}+1} \frac{2}{k} \sum_{j=1}^{k} \frac{1}{j} - \left( \frac{H_{\epsilon_{NC}+1}}{\lambda} \right)^2 \\
&= \frac{1}{\lambda^2} \cdot \left( \sum_{k=1}^{\epsilon_{NC}+1} \sum_{j=1}^{k} \frac{2}{jk} - \sum_{k=1}^{\epsilon_{NC}+1} \sum_{j=1}^{\epsilon_{NC}+1} \frac{1}{jk} \right)
\end{aligned} \tag{7}$$

Using the fact that the sum $\sum_{k=1}^{\epsilon_{NC}+1} \sum_{j=1}^{\epsilon_{NC}+1} \frac{1}{jk}$ can be split into two parts for $j = k$ and $j \neq k$ gives:

$$\sum_{k=1}^{\epsilon_{NC}+1} \sum_{j=1}^{\epsilon_{NC}+1} \frac{1}{jk} = \sum_{k=1}^{\epsilon_{NC}+1} \frac{1}{k^2} + \sum_{k=1}^{\epsilon_{NC}+1} \sum_{j=1}^{k-1} \frac{2}{jk}$$

where the factor of 2 in the second term appears due to symmetry of indices. Thus, substituting for $\sum_{k=1}^{\epsilon_{NC}+1}\sum_{j=1}^{k}\frac{2}{jk}$ in eq. (7) gives rise to $\sum_{k=1}^{\epsilon_{NC}+1}\sum_{j=1}^{\epsilon_{NC}+1}\frac{1}{jk}-\sum_{k=1}^{\epsilon_{NC}+1}\frac{1}{k^2}+\sum_{k=1}^{\epsilon_{NC}+1}\frac{2}{k^2}$, where summing $j$ from 1 to $k-1$ gives rise to the first two terms and the last term appears when $j=k$. Therefore,

$$\begin{aligned}\mathrm{Var}(F_{\mathrm{peak}}) &= \frac{1}{\lambda^2}\cdot\left(\sum_{k=1}^{\epsilon_{NC}+1}\sum_{j=1}^{\epsilon_{NC}+1}\frac{1}{jk}-\sum_{k=1}^{\epsilon_{NC}+1}\frac{1}{k^2}+\sum_{k=1}^{\epsilon_{NC}+1}\frac{2}{k^2}-\sum_{k=1}^{\epsilon_{NC}+1}\sum_{j=1}^{\epsilon_{NC}+1}\frac{1}{jk}\right)\\ &= \frac{1}{\lambda^2}\cdot\sum_{k=1}^{\epsilon_{NC}+1}\frac{1}{k^2}\end{aligned} \tag{8}$$

As expected, these expressions for the mean and variance agree with the literature on order statistics of exponential distributions: they are the mean and the variance of the maximum out of $\epsilon_{NC}+1$ exponentially drawn values [4].

#### S1.3 Scaling of minimum evolvability $\bar{\epsilon}_{NC}$ required for navigability

Navigability is a continuous quantity characterising a GP map: it captures the probability that there is at least one fitness-increasing (i.e. accessible) path from a randomly chosen initial phenotype to a target phenotype whose fitness is set to one [6]. Here, we only focus on distinguishing non-navigable GP maps (with navigability $\approx 0$) from potentially navigable ones. Since accessible paths can only exist on a GP map if they exist on the corresponding NC graph, we will focus on the number of accessible paths on a NC graph.

To simplify the calculations, we will assume that the GP map has one NC per phenotype. Then, each NC has a unique fitness value and its degree equals its evolvability. Further, we assume that the NC graph's degree distribution is narrow and can be characterised by a single evolvability value $\epsilon_{NC}$. With these two simplifications, we estimate the number of accessible $k$-step paths on the NC graph (adapting similar calculations for abstract directed acyclic graphs [7], but accounting for the source fitness):

1. **Combinatorics of fitness-increasing length-$k$ sequences of NCs:** A length-$k$ path involves the source, target and $k-1$ intermediate nodes. If the NC graph was a *complete graph* (i.e. with an edge between any two nodes), then these $k-1$ intermediate nodes could be any of the $n_p-2$ nodes of the graph other than the source or target. Since these $k-1$ nodes can only be traversed in order of increasing fitness, their order is fixed, and there are $\binom{n_p-2}{k-1}$ fitness-increasing sequences of intermediate nodes.

2. **Connectivity of the NC graph and source fitness**: Next, we correct for the fact that the NC graph is not complete and that some NCs are lower in fitness than the source and thus cannot appear as intermediate nodes.

   - For each step on the path, we approximate the probability that this step is mutationally possible by $\frac{\epsilon_{NC}}{n_p-1}$ since each node is connected to $\epsilon_{NC}$ out of $n_p-1$ NCs.
   - Further, only NCs fitter than the source can appear as intermediate nodes. If we assume a uniform fitness distribution[1], and the source has a fitness $1-\beta$, then this is the case with probability $\beta^{k-1}$

[1]Since accessible paths only depend on fitness rankings, the result should not depend on the fitness distribution as long as it remains continuous.

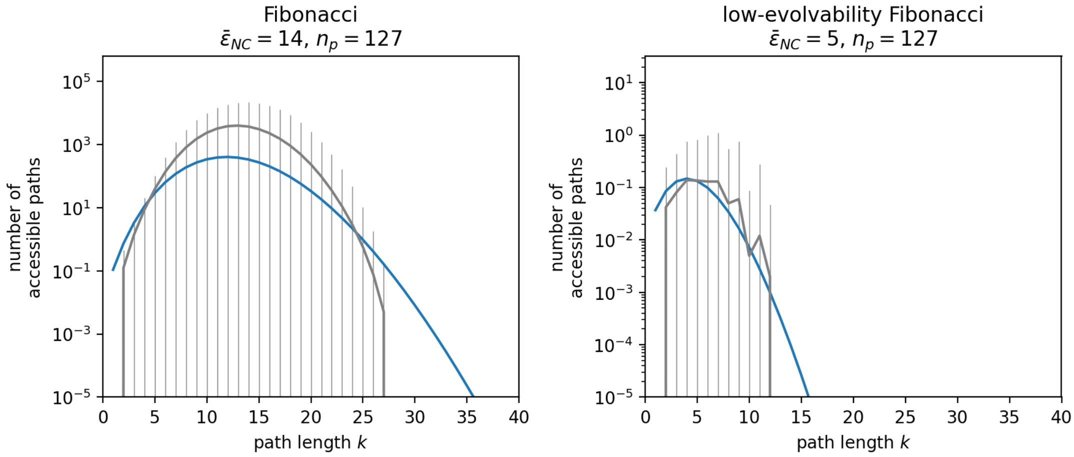

Figure S1: **Average number of accessible paths of length $k$ in the NC graph of the Fibonacci model and its low-evolvability version:** here, we test an intermediate result of our navigability calculations (eq. (9), blue line): the number of accessible $k$-step paths from a source to a target on a NC graph. The grey curves represent simulation averages (with standard deviations). We see clear differences between the Fibonacci GP map on the left and its low-evolvability version on the right (note the different y-axis scales). These differences are well-approximated by the estimate from eq. (9), based only on two GP map parameters: $\bar{\epsilon}_{NC}$ and $n_p$. Parameters: average taken over $10^2$ PF maps & 10 source-target pairs for each PF map; sequence length $L = 7$ & alphabet size $K = 3$ (note that this GP map is very small for reasons of computational feasibility).

Thus, we approximate the number of accessible k-step paths as:

$$\langle \text{number of k-step paths} \rangle(\beta) = \binom{n_p - 2}{k-1} \left( \frac{\epsilon_{NC}}{n_p - 1} \right)^k \beta^{k-1}$$

Since we are interested in the *average* number of paths over all PF maps, we first take the mean over $\beta$, which characterises the source fitness ($F_{\text{source}} = 1 - \beta$) and is uniformly distributed between zero and one:

$$\begin{aligned} \langle \text{number of k-step paths} \rangle &= \binom{n_p - 2}{k-1} \left( \frac{\epsilon_{NC}}{n_p - 1} \right)^k \int_0^1 d\beta \cdot \beta^{k-1} \\ &= \binom{n_p - 2}{k-1} \left( \frac{\epsilon_{NC}}{n_p - 1} \right)^k \times \frac{1}{k} \end{aligned} \tag{9}$$

To test this intermediate result against simulation data, we applied it to the Fibonacci model and its low-evolvability version (Fig. S1). For this, we need to find a single representative $\epsilon_{NC}$ describing the full distribution of $\epsilon_{NC}$ on each NC graph, and used the geometric mean $\bar{\epsilon}_{NC}$ of all $\epsilon_{NC}$ values (since $\epsilon_{NC}$ enters the calculation as a product, with the single zero-evolvability NC in the low-evolvability version treated as 0.01). We find that the simple estimate from eq. (9) approximates the order of magnitude, as well as the contrast between the Fibonacci model and its low-evolvability version well despite this oversimplified treatment of the degree distribution.

Having approximated the number of length-$k$ accessible paths on the NC graph, we can sum over $k$ to estimate the total number of paths:

$$\langle \text{number of paths} \rangle = \sum_{k=1}^{n_p-1} \binom{n_p-2}{k-1} \left(\frac{\epsilon_{NC}}{n_p-1}\right)^k \times \frac{1}{k}$$

For simplicity, let us denote $N = n_p - 1$ and $c = \frac{\epsilon_{NC}}{n_p-1}$. Then we have

$$\langle \text{number of paths} \rangle = \sum_{k=1}^{N} \binom{N-1}{k-1} c^k \times \frac{1}{k}$$

Using the binomial identity $\binom{N-1}{k-1} = \frac{k}{N}\binom{N}{k}$:

$$\begin{aligned} \langle \text{number of paths} \rangle &= \frac{1}{N} \sum_{k=1}^{N} \binom{N}{k} c^k \\ &= \frac{1}{N}\left((1+c)^N - 1\right) \end{aligned} \tag{10}$$

Note that eq. (10) estimates the mean number of accessible paths *on the NC graph.* While the number of paths in genotype space can be much higher due to neutral mutations, accessible paths in genotype space can only exist if at least one accessible path exists on the NC graph. Thus, we can use the fact that the navigability is bounded by $P(\text{at least one path exists}) \leq \langle \text{number of paths} \rangle$ [8], to find that a NC graph will be non-navigable (navigability $< 0.1$) if:

$$0.1 > \frac{(c+1)^N - 1}{N}$$

Since we are interested in the case $n_p \gg 1$, we have $N \approx n_p$ and $1/N \ll 1$. Then we can simplify to:

$$\begin{aligned} 0.1 &> \frac{(c+1)^{n_p}}{n_p} \\ c &< \left(\frac{n_p}{10}\right)^{\frac{1}{n_p}} - 1 \end{aligned}$$

And thus, substituting $c = \frac{\epsilon_{NC}}{n_p-1} \approx \frac{\epsilon_{NC}}{n_p}$:

$$\epsilon_{NC} < n_p \left(\left(\frac{n_p}{10}\right)^{\frac{1}{n_p}} - 1\right) \tag{11}$$

Thus, low-evolvability GP maps, with evolvabilities below the bound in eq. (11) are expected to have low navigability.

An interesting question is whether the reverse statement also holds, i.e. that high-evolvability GP maps have high navigability. To put such a lower bound on navigability, we would need to know not only the expected number of paths, but also its variance for the following reason [8]: NC graphs with a high expected number of paths could nevertheless be of low navigability if there were no paths for most PF maps and

$\gg 1$ paths for a small number of PF maps. Computing this variance is an interesting topic for future work (similar to work on tree graphs [9]). However, even without computing the variance, it is clear that a GP map will become navigable in the maximum-evolvability limit $\epsilon \to n_p$, since single-step paths from a source to a target become possible and these are always accessible.

It is important to emphasise the caveats in this derivation, especially: (1) It assumes that the network is characterised by a single degree $\epsilon_{NC}$, which is not true for real NC graphs. In applications of eq. (11) (see section S2.4), we will take the geometric mean $\bar{\epsilon}_{NC}$ as a proxy[2]. (2) The calculations assume that any NC is equally likely to be connected to any other NC, but real NCs may be more likely to be connected to other NCs that are close in genotype space (although some NCs can span the maximum genotypic distance in the GP map [10]). (3) The combinatoric terms assume a single NC per phenotype. (4) The fitness of the source phenotype is accounted for in a highly simplified way via the parameter $\beta$.

Due to these caveats, the most important takeaway from this calculation may not be the exact quantitative bound in eq. (11), but the qualitative result that low-evolvability NC graphs are non-navigable, as hypothesised in [11], and that this bound also depends on combinatoric terms, here the number of phenotypes $n_p$.

#### S1.4 Analytic expression for ruggedness in the Fibonacci model

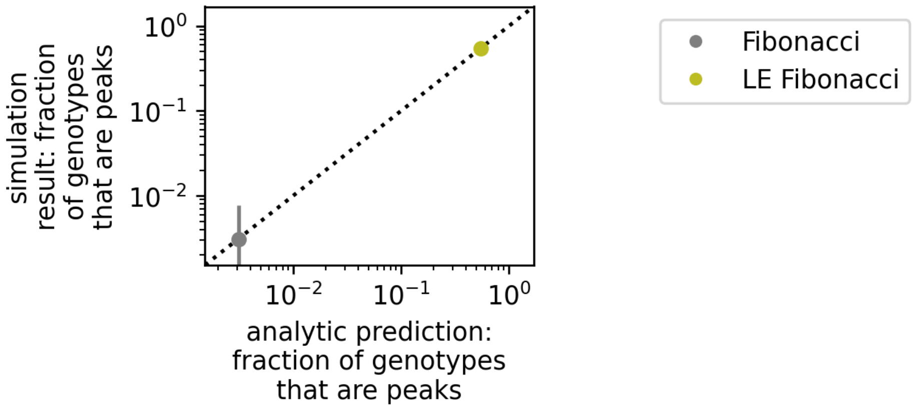

Figure S2: **Analytic ruggedness calculations for the Fibonacci model:** in the Fibonacci model, NC evolvabilities $\epsilon_{NC}$ and NC sizes $|NC|$ can be computed analytically, meaning that the ruggedness can be predicted purely analytically (eq. (14)), without relying on any computationally derived GP map properties. Here, simulation data (y-axis) is plotted against the purely analytic prediction (x-axis, eq. (14)). Parameters: sequence length $L = 12$, alphabet size $K = 3$, data based on $10^3$ sampled PF maps.

One advantage of the Fibonacci model is that its GP map characteristics, such as evolvability, can be calculated analytically as parametric equations of a single parameter $l$, the number of sites before the first stop codon [12][3]. Since each phenotype corresponds to a single NC in the Fibonacci model, NC evolvabilities and sizes are equivalent to phenotypic evolvabilities and sizes, which are given by the following parametric

[2]Since a single zero-evolvability node turns the geometric mean to zero, we treated zero evolvability values as 0.01

[3]To simplify expressions, we use a slightly different notation than the original paper, which included the stop codon in $l$ [12].

equations [12]:

$$|NC|(l) = K^{L-l-1} \tag{12}$$

Similarly, we can write down the evolvability as [12][4]

$$\epsilon_{NC}(l) = (K-1) \times l + (K-1) \times \left( \sum_{i=0}^{L-l-2} (K-1)^i \right) \tag{13}$$

There are $S$ NCs with evolvability $\epsilon_{NC}(l)$ and size $|NC|(l)$, where $S(l)$ is given by [12]:

$$S(l) = (K-1)^l$$

In the low-evolvability reference model, the sizes $|NC|$ and numbers $S$ of NCs are unchanged; only the evolvability has to be re-calculated. Since in our version of the low-evolvability model, we only allow (K-2) mutations per site before the stop codon, the evolvability in the low-evolvability model is $\epsilon_{NC}(l) = (K-2) \times l$, rather than $(K-1) \times l$ as in the original formulation of the model [12].

With this, we can compute the ruggedness analytically rather than relying on the computational data for evolvabilities and NC sizes:

$$\langle \text{fraction of genotypes in peaks} \rangle = \frac{1}{K^L} \sum_{l=0}^{L-1} \frac{S(l) \times |NC|(l)}{\epsilon_{NC}(l) + 1} \tag{14}$$

This analytic prediction is shown in Fig. S2 and is in excellent agreement with the simulation data for both the standard and low-evolvability Fibonacci models.

#### S1.5 Extrapolating ruggedness to large GP maps

The sequence lengths of the GP maps studied in this paper are limited by computational feasibility. More realistic GP maps would have longer sequence lengths $L$, and thus a higher number of genotypes $N_G = K^L$, a higher number of phenotypes $n_p$ and a higher number of NCs $n_c$. To estimate how ruggedness might scale if we studied larger GP maps, we need to make analytic extrapolations.

##### S1.5.1 Scaling in HoC models as a baseline

As a baseline for our extrapolations, we take a classic fitness landscape baseline, the "House-of-cards (HoC)" landscape [1, 13], where each genotype maps to a random fitness value. Since each geotype's fitness corresponds to an independent draw from a continous distribution, genotypes do not have neutral neighbours and form single-genotype NCs. Since each single-genotype NC has $(K-1)L$ mutational neighbours, we can define a 'HoC-evolvability' of $\epsilon_{\text{HoC}} = (K-1)L$ distinct fitness values around each NC, giving a ruggedness of $R_{\text{HoC}} = \frac{1}{(K-1)L+1} \approx \frac{1}{(K-1)L}$ [1]. We could also introduce deleterious genotypes into the HoC model, to build a more accurate baseline for our GP maps. This would reduce the 'HoC evolvability' by a factor depending on the fraction of deleterious genotypes and increase the HoC ruggedness.

[4]Here we ignore mutations to the 'undefined'/deleterious phenotype with no stop codon.

##### S1.5.2 Scaling in GP maps

The analogous calculation for GPF maps is more complex because genotypes mapping to the same phenotype do form large connected neutral components (NCs) [14]. Thus, we cannot treat each genotype separately and need to estimate how NC sizes and evolvabilities scale with $L$. Here we will work with a normalised version of a NC's size, the *NC frequency* $f_r$, which equals the size of a neutral component, normalised by the total number of genotypes in the GP map $f_r = |NC_r|/K^L$. The index $r$ denotes the rank, i.e. the number of NCs with larger or equal size (see Fig. S3).

**S1.5.2.1 NC frequency scaling** Since the distribution of phenotypic frequencies is highly biased in GP maps [15–17], we also expect the NC frequency distribution to be biased. Based on our computational data in Fig. S3, we will work with a simple power-law ansatz:

$$f_r = A/r^m \tag{15}$$

Here, the rank $r$, the number of NCs of equal or higher size, can go from $r = 1$ for the largest NC to $r = n_c$ for the smallest NC (where $n_c$ is the total number of NCs in the GP map).
The use of power laws is also supported by theoretical arguments based on constrained and unconstrained sites [18], which suggest a power law for *phenotypic* frequencies, which may translate to a power law in *NC* frequencies. Note, however, that these arguments can also predict other functional forms [19], including the log-normal/binomial distributions found computationally for the RNA secondary structure GP map [20].

The parameter $A$ in eq. (15) follows from the normalisation $1 = \sum_r f_r \approx \int_{r=1}^{r=n_c} f_r dr$:

$$\begin{aligned} A &= \frac{(m-1)}{1-n_c^{(1-m)}} && \text{if } m \neq 1, \\ A &= \frac{1}{\ln(n_c)} && \text{if } m = 1. \end{aligned}$$

For simplicity we will assume for $m \neq 1$ that $n_c$ is large (meaning $\ln n_c \gg 1/|m-1|$). Then we have:

$$\begin{aligned} A &\approx \frac{1-m}{n_c^{(1-m)}} && \text{if } m < 1, \\ A &\approx m-1 && \text{if } m > 1. \end{aligned}$$

**S1.5.2.2 Power-law frequency distributions capture a range of phenomenologies** To illustrate the diverse range of phenomenologies encompassed by the power-law frequency distributions for different $m$, we will show how they differ in key characteristics.

The first characteristic we will compare is, how many NCs are larger or equal than the average frequency $\langle f_r \rangle = 1/n_c$. This is given by $r_{\text{ave}}$, the rank at which $f_{r_{\text{ave}}} = 1/n_c$.

The second characteristic we will compare is $r_x$, the number of largest NCs required to account for a fraction $0 < x \leq 1$ of all genotypes. This is computed by taking the integral $x = \int_{r=1}^{r=r_x} f_r dr$ to give:

$$\begin{aligned} r_x &= \left(1 - x + x n_c^{-(m-1)}\right)^{-1/(m-1)} && \text{if } m \neq 1, \\ r_x &= n_c^x && \text{if } m = 1. \end{aligned}$$

These quantities highlight the differences between power laws with $m = 1$, $m > 1$, and $m < 1$ (in the limit of large GP maps with large $L$, $n_p$ and $n_c$):

- For the pure Zipfian case $m = 1$:

$$r_{50\%} = n_c^{1/2}, \quad r_{\text{ave}} = \frac{n_c}{\ln(n_c)} \ll n_c, \quad \text{the smallest } f_{n_c} = \frac{1}{\ln(n_c)n_c} \ll \langle f_r \rangle = \frac{1}{n_c}.$$

  Only a small minority of the largest NCs account for half of all genotypes, and only a small minority of NCs are above-average in size.

- For the "super-Zipfian" case $m > 1$:

$$r_{50\%} \approx 2^{1/(m-1)}, \quad r_{\text{ave}} = (m-1)^{1/m} \cdot n_c^{1/m} \ll n_c, \quad f_{n_c} \approx \frac{m-1}{n_c^m} \ll \langle f_r \rangle$$

  To leading order in $n_c$, the number of large NCs accounting for half the genotypes, $r_{50\%}$, is independent of the number of NCs, and thus of the size of the GP map. Since the smallest NC must contain at least one genotype, this scaling is only valid for $(N_G/n_c^m) \geq 1/(m-1)$.

- For the "sub-Zipfian" case $0 < m < 1$:

$$r_{50\%} \approx 2^{-1/(1-m)} n_c \propto n_c, \quad r_{\text{ave}} = (1-m)^{1/m} n_c \propto n_c, \quad f_{n_c} = \frac{1-m}{n_c} \propto \langle f_r \rangle$$

  In contrast to the $m \geq 1$ case, for the sub-Zipfian case, both $r_{50\%}$ and $r_{\text{ave}}$ are directly proportional to $n_c$ and the smallest frequency is on the same order of magnitude as the average NC frequency.

These characteristics illustrate that the three options for the exponent $m$ define a diverse set of GP maps. Thus, as a set they may cover key aspects of the spectrum of scaling behaviour for a wider range of GP maps, even though the exact frequency distributions of these GP maps are likely to be more complex than simple power-law forms.

**S1.5.2.3 NC evolvability scaling** To approximate the ruggedness, we also need to estimate the evolvability $\epsilon_r$ of the $r$th NC. Here, we make the ansatz that evolvability scales as a power law with frequency $\epsilon_r \propto f_r^\beta$, based on the relationships seen in our GP maps (with $0 \leq \beta \leq 1$, Fig. S4). To bound the normalisation constant, we note that the evolvability cannot exceed the total number of phenotypes $n_p$. Thus, we will write the highest evolvability corresponding to the largest NC (with rank $r = 1$) as $\epsilon_1 = \alpha n_p$, where $\alpha \leq 1$ (for example, in one of the largest exhaustively studied GP maps, that of $L = 20$ RNA secondary structures, $\alpha > 0.75$ [21]).

Assuming that the power law scaling approximately holds for the full range of frequencies, without saturation effects at low or high frequencies (note that this is an approximation, for example in the Fibonacci model in Fig S4), we have the ansatz:

$$\epsilon_r \;=\; \alpha n_p \left(\frac{f_r}{f_1}\right)^\beta \tag{16}$$

In this scaling, the largest NC has an evolvability of $\epsilon_1 = \alpha n_p$, and the smallest NC (of rank $n_c$) an evolvability of $\epsilon_{\min} = \alpha n_p n_c^{-\beta m}$.

**S1.5.2.4 Combining frequencies and evolvabilities to compute ruggedness** Our scalings for the frequency distribution and evolvability, together with the fact that $f_r = |NC_r|/K^L$ by definition, allow us to rewrite eq. (2) for ruggedness as

$$\begin{aligned}
\langle \text{fraction of genotypes in peaks} \rangle = R &= \sum_{r=1}^{n_c} \frac{f_r}{\epsilon_r + 1} \\
&\approx \frac{f_1^{\beta}}{\alpha n_p} \sum_{r=1}^{n_c} f_r^{1-\beta} \\
&= \frac{A}{\alpha n_p} \int_{r=1}^{r=n_c} \frac{1}{r^{m(1-\beta)}}\, dr \\
&= \frac{A}{1-m(1-\beta)} \left[ \frac{n_c^{m\beta}}{\alpha n_p} n_c^{(1-m)} - \frac{1}{\alpha n_p} \right] \quad \text{if } m(1-\beta) \neq 1 \\
&= \frac{A}{(1-m(1-\beta))\epsilon_{\min}} \left[ n_c^{(1-m)} - n_c^{-\beta m} \right] \qquad (17)
\end{aligned}$$

In the limit of large GP maps ($n_c \to \infty$) we find the following scaling:

- For $m = 1$, the expression in the square brackets is dominated by the first term, so the dominant scaling is $R \approx \frac{1}{\beta \ln n_c} \frac{1}{\epsilon_{\min}} \ll \frac{1}{\epsilon_{min}}$
- For $m > 1$, there are two cases:[5] First, $m - 1 < m\beta$, or equivalently $m < 1/(1-\beta)$ the dominant term is $\propto \frac{1}{n_c^{(m-1)} \epsilon_{\min}}$. If $m > 1/(1-\beta)$ (the large $m$ limit) then ruggedness is dominated by the evolvability of the largest NC ($\epsilon_1 = \alpha n_p$) and $R \propto \frac{1}{n_c^{m\beta} \epsilon_{\min}}$. In either case, for large $n_c$, we have $R \ll \frac{1}{\epsilon_{min}}$.
- For $m < 1$ the dominant scaling is $R \approx \frac{1-m}{1-m(1-\beta)} \frac{1}{\epsilon_{\min}} \sim \mathcal{O}(\frac{1}{\epsilon_{\min}})$

To make further progress on our scaling arguments for the ruggedness, we need to make some assumptions about the minimum evolvability in the map, $\epsilon_{min}$.

**S1.5.2.5 Scaling assuming the minimum evolvability $\epsilon_{\mathbf{min}} \approx \epsilon_{\mathbf{HoC}}$** Let us first start with a simple assumption about $\epsilon_{min}$, namely that it is independent of the size of the smallest NC and that it scales like the evolvability of a single genotype in the HoC model, $\epsilon_{\min} \propto \epsilon_{\text{HoC}} \sim 1/L$. On the one hand, this may be an underestimate of the true minimal evolvability since the smallest NC may have many genotypes, and so many more mutational neighbours than a single-genotype NC in the HoC model. On the other hand, the GP map evolvability needs to account for repeated phenotypes in a mutational neighbourhood, which reduce the evolvability compared to the HoC model: repeated phenotypes can occur by chance, depending on the phenotypic frequency distribution, or due to genetic correlations, i.e. the clustering of genotypes mapping to the same phenotype present in realistic GP maps [14].

[5] Note that for $m > 1$, the power-law frequency distribution is only valid if the smallest NC contains at least one genotype, giving the condition $n_c^{m-1}/(m-1) \leq n_G/n_c$ (see section “Power-law frequency distributions capture a range of phenomenologies"). This limits $m$ to a value that depends on the number of genotypes $N_G$ and the number of NCs $n_c$.

With this assumption that $\epsilon_{\text{min}} \approx \epsilon_{\text{HoC}}$, the ruggedness $R$ of a GP map with $m \geq 1$ will be much smaller than $1/\epsilon_{\text{min}}$, and thus much smaller than that of the corresponding HoC baseline. This small ruggedness is due to the fact that larger, high-evolvability NCs take up such a large fraction of the genotype space for $m \geq 1$ that the smallest, low-evolvability NCs do not contribute to the dominant scaling in the ruggedness calculations.

For $m < 1$, on the other hand, the assumption that $\epsilon_{\text{min}} \approx \epsilon_{\text{HoC}}$ leads to a ruggedness that scales like the ruggedness in the HoC model. However, for $m < 1$ the assumption that $\epsilon_{\text{min}} \approx \epsilon_{\text{HoC}}$ is most likely to be an underestimate since even the smallest NC is relatively large, on the same order of magnitude as the mean NC size (see section "Power-law frequency distributions capture a range of phenomenologies") .

**S1.5.2.6 Scaling when the minimum evolvability $\epsilon_{\text{min}}$ grows with the size of the smallest NC** Let us revisit the $m < 1$ case, where even the smallest NC is on the same order of magnitude as the mean NC size (see section "Power-law frequency distributions capture a range of phenomenologies"), and thus likely to have a higher evolvability than single-genotype NCs in the HoC model. For example, we might assume that a NC made up of $x$ genotypes should have access to a $x$-times as many distinct phenotypes than a NC made up of a single genotype in the HoC model. This would be an overestimate because it ignores repeated phenotypes, either due to diminishing returns (if the same high-frequency phenotypes appear in the neighbourhood repeatedly due to phenotypic bias [22]) or due to genetic correlations [14]. Thus, the scaling might be non-linear, as in our initial evolvability scaling, where frequency differences *within* a GP map are related to evolvability differences *within* a GP map via a power law with exponent $\beta < 1$ (see eq. (16)). However, we do not have to assume a specific functional form: we simply assume that $\epsilon_{min} \gg \epsilon_{HoC}$ as long as the minimum NC size is much larger than one, i.e. $K^L f_{n_c} \gg 1$. With this assumption, the ruggedness, which scales as $\epsilon_{\text{min}}^{-1}$ for the $m < 1$ case, will be much lower than the HoC ruggedness, i.e. $R \ll R_{\text{HoC}}$.

To sum, up, we get the following intuitive result: if even the smallest NCs are large and have much higher evolvabilities than a single-genotype NC in the HoC model, then the ruggedness is much smaller than in the HoC expectation even for the $m < 1$ power laws.

**S1.5.2.7 Discussion: testing extrapolations through sampling** The big question then is whether these extrapolations will hold for realistic, biophysical GP maps in the long-$L$ limit. Here some caution is needed. While the power-law assumption permits a broad range of qualitatively different frequency distributions (see section "Power-law frequency distributions capture a range of phenomenologies"), it needs to be tested against GP maps in the large-$L$ limit. For this, we cannot simply rely on scaling laws for phenotype frequencies since the genotypes mapping to a single phenotype (a 'neutral set') can fragment into disconnected neutral components for a variety of reasons, including biophysical reasons such as base pairing in the RNA secondary structure GP map [23]. Similarly, our power-law assumption for the scaling of evolvabilities with frequencies, as well as our assumptions about minimum evolvabilities, need to be tested in large-$L$ GP maps. Since exhaustive analysis will not be feasible for large-$L$ GP maps, frequencies and evolvabilities will have to be estimated using sampling methods. Such analyses have successfully revealed the scaling of *phenotypic* quantities for RNA secondary structures [20] (such as neutral set sizes) and a sampling method for NC *sizes* exists [24]. However, more work is needed to develop accurate sampling approaches to estimate NC *evolvabilities* [25].

| GP map | number NCs | number genotypes | number phenotypes | number zero-evolv. NCs | mean size zero-evolv. NCs | prevalence deleterious phenotype |
|---|---|---|---|---|---|---|
| HP_20 | 6,586 | 1,048,576 | 5,310 | 2,067 | 2 | 98 % |
| HP3x3x3s | 732,157 | 134,217,728 | 49,807 | 151,640 | 1 | 94 % |
| HP5x5s | 6,785 | 33,554,432 | 549 | 586 | 1 | 82 % |
| HP25 | 148,254 | 33,554,432 | 107,336 | 36,872 | 2 | 98 % |
| RNA structure | 645 | 16,777,216 | 57 | 0 | | 85 % |
| self-assembly | 1,347 | 16,777,216 | 13 | 0 | | 54 % |
| RNA-binding | 839 | 16,384 | 305 | 5 | 1 | 87 % |
| Fibonacci | 4,095 | 531,441 | 4,095 | 0 | | 1 % |
| LE Fibonacci | 4,095 | 531,441 | 4,095 | 1 | 177,147 | 1 % |

Table 1: **Characteristics of the GP map models used throughout this paper:** for each GP map, the table gives the number of NCs, the number of genotypes $N_G = K^L$ (which includes those folding to the 'deleterious' phenotype), the number of distinct viable phenotypes, the number of isolated NCs with $\epsilon_{NC} = 0$, the mean size of isolated NCs and the prevalence of the 'deleterious' phenotype among all genotypes. The biophysical GP maps are constructed from Greenbury et al.'s [6] datasets (several characteristics are also reported in ref [6], and included here for completeness), and are based on the following models: the HP lattice models of protein tertiary structure [26, 27], the Polyomino self-assembly tile model of protein quaternary structure (proposed in this form in [28], but based on a longer history of tile-assembly models, see [29]) and a free-energy-based RNA secondary structure folding model implemented in the ViennaRNA package [30]. The data underlying the RNA-binding map was curated by Payne and Wagner [31] from the CISBP-RNA database, and we constructed a categorical, many-to-one GP map from this data as described in the text. Finally, we have the Fibonacci model and its low-evolvability version (adapted from the 'gene-like model' and the corresponding reference model [12], which is a generalisation of the original 'Fibonacci' model [32]), with a minor adjustment to guarantee symmetry of allowed mutations as described in the main text.

### S2 Supplementary computational analyses

#### S2.1 Additional NC graph analyses

##### S2.1.1 Number, evolvabilities and sizes of NCs

In our ruggedness predictions, the number, evolvabilities, and sizes of NCs are crucial. In Table 1, we show the number of NCs and further key quantities for each GP map from the main text. We also show the size distribution of NCs in Fig. S3, and the relationship between NC size and NC evolvability in Fig. S4.

As expected from prior literature, see e.g. [23, 33], we find that large NCs tend to be more evolvable in all GP maps with the exception of the low-evolvability Fibonacci model, where evolvability is low for all phenotypes by construction and decreases with NC size [12]. We further note that, out of all biophysical GP maps, the self-assembly GP map has the flattest slope in the NC-size-evolvability plot, indicating that even large NCs only reach moderate evolvabilities. Thus, even large NCs have a comparatively high likelihood of being peaks and the ruggedness (Fig 2 in the main text) is comparatively high in this map. However, this feature may be an artefact of the limited size of the model in particular [6]: with a sequence length of $L = 8$,

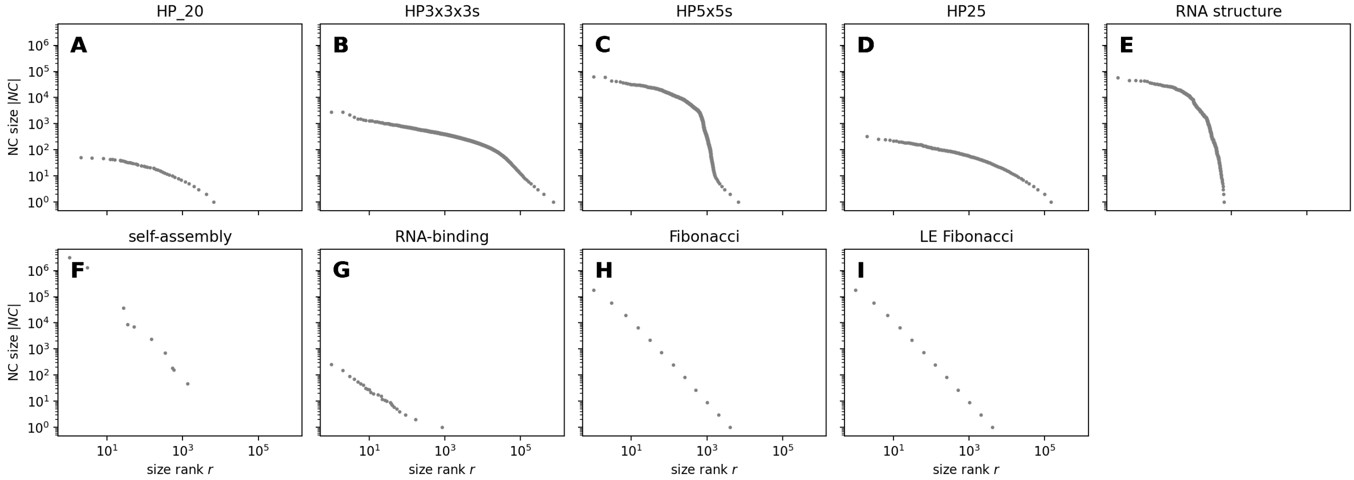

Figure S3: **Distribution of NC sizes $|NC|$:** in the scaling calculations (section S1.5), the distribution of NC sizes is important. Thus, for each GP map, we take each NC and plot its NC size $|NC|$ against the NC size rank, i.e. the number of NCs with equal or higher size.

only 13 distinct viable phenotypes exist, and therefore evolvabilities have to be integers between $\epsilon_{NC} = 0$ and $\epsilon_{NC} = 12$. The number of phenotypes is higher for longer sequences and larger alphabets [28], but these GP maps are difficult to store and analyse exhaustively because the number of genotypes grows rapidly with sequence length as $K^L$, especially for the large alphabets typical of the self-assembly models.

Moreover, we note that some GP maps have completely isolated NCs with $\epsilon_{NC} = 0$ (i.e. NCs with mutational connections to the deleterious phenotype). These isolated peaks are particularly common in the HP protein model, and constitute one of the differences between this model and the RNA secondary structure model (see [34, 35] for a more detailed discussion on qualitative and quantitative differences). Such isolated NCs are guaranteed to be peaks under any fitness assignment. Since such isolated peaks may never be reached by evolving populations, we repeat our ruggedness analysis with isolated peaks excluded in section S2.2.2.

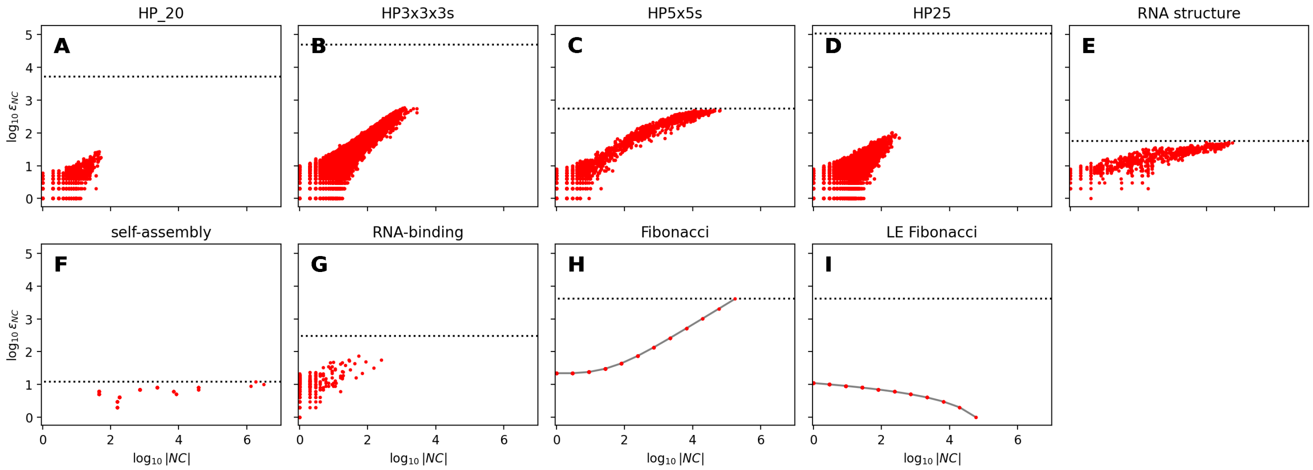

Figure S4: **Large NCs with high $|NC|$ tend to be more evolvable (higher $\epsilon_{NC}$), except in the 'low-evolvability' Fibonacci map.** The relationship between NC size and evolvability is important for the prevalence of peaks in genotype space. Thus, for each GP map, we take each NC and plot the logarithm of the NC evolvability $\log_{10} \epsilon_{NC}$ against the logarithm of the NC size $\log_{10} |NC|$ (excluding zero-evolvability NCs values due to the log scale). In addition to the computational data (red), analytic relationships are shown for the Fibonacci models (grey, eqs (12-13)). The maximum possible evolvability $n_p - 1$ (where $n_p$ is the number of phenotypes in the map) is drawn as a black dotted line.

##### S2.1.2 Treatment of evolvabilities for main text Fig.4

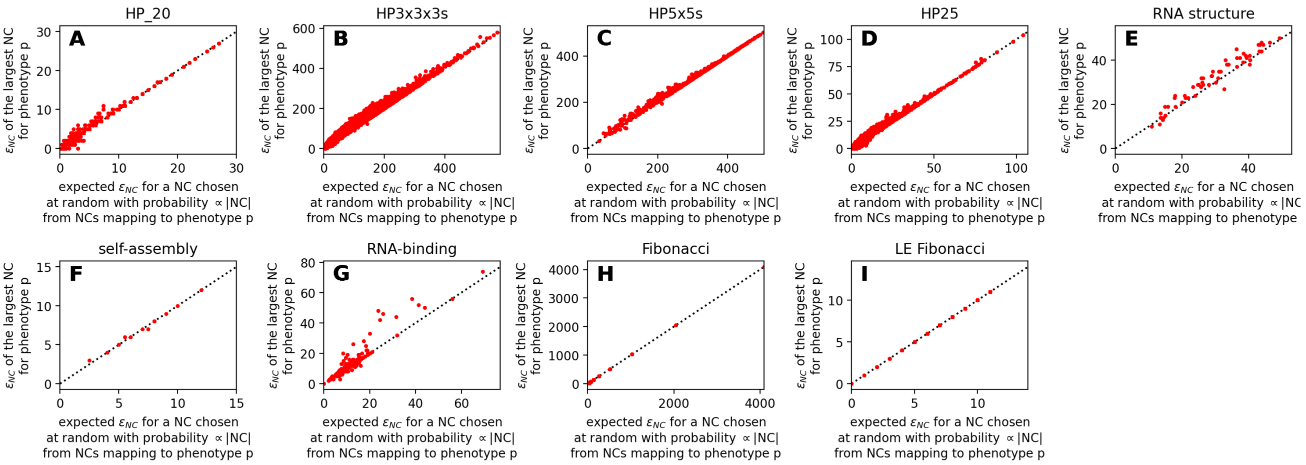

Figure S5: $\epsilon_{NC}$ **of the largest NC mapping to a given phenotype vs. expected** $\epsilon_{NC}$ **when a NC for the same phenotype is randomly selected with a probability proportional to its size:** we find that the two evolvabilities are of similar orders of magnitude in our maps.

In Fig. 4 in the main text, all navigability simulations were based on the full NC graph, but to obtain a single evolvability value representing each NC graph, several simplifications were made. First, we simplified NC graphs to versions containing a single NC per phenotype, and then we summarised the evolvabilities in these simplified graphs by their geometric mean. Here, we investigate both simplifications.

First, we turn to one of the NC graph simplifications, which only retains the largest NC for every phenotype. Specifically, we focus on the impact this may have for the initial condition: in the navigability simulations, the initial NC is chosen from the 'neutral set' of all NCs mapping to the source phenotype with a probability proportional to the NC size. Thus, it is possible that the initial NC will be a small, low-evolvability NC from the neutral set, which may be an evolutionary dead end and is not accounted for in the simplified graph. To investigate this potential discrepancy, we took the neutral set of each phenotype in each GP map, and compared the evolvability of the largest NC in the neutral set to the expected evolvability of a NC when sampling from NCs in the neutral set with a probability proportional to the NC sizes (Fig. S5). We find that the two evolvabilities are of similar orders of magnitude in our GP maps.

Secondly, we turn to the second step, after the NC graph has been simplified, and compare several ways of representing a NC graph's evolvability distribution by a single typical evolvability: one could take the arithmetic mean or the median of all evolvabilities, but we chose to use the geometric mean instead since evolvabilities enter the calculation as a product (see section S1.3). Fig. S6 illustrates that the order of magnitude of the typical evolvability, which is key in our navigability analysis, does not depend strongly on this choice.

Of course, these two tests only cover specific aspects of our simplifications and many caveats remain.

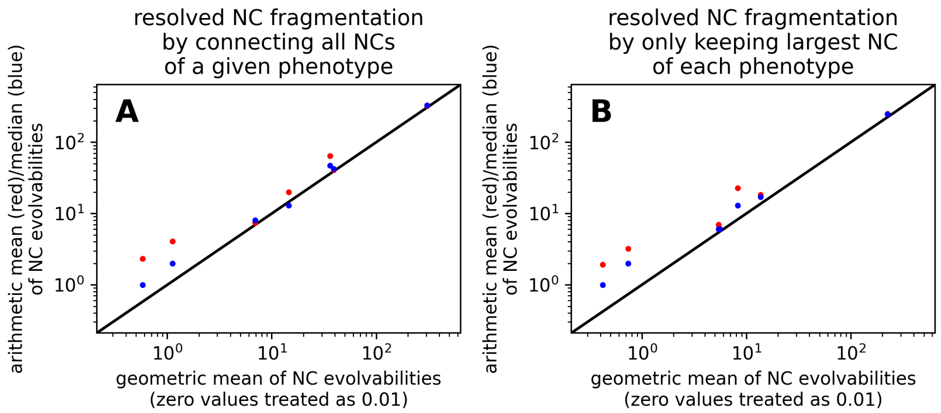

Figure S6: **Comparisons of the median and arithmetic/geometric mean evolvability in a NC graph:** In Fig. 4 in the main text, we use the geometric mean to get a representative evolvability value for a NC graph (with zero values treated as 0.01, such that a single zero-evolvability NC does not set the mean to zero). Here, we compare this geometric mean (x-axis) to the arithmetic mean (red) and the median (blue) for the GP maps included in Fig. 4. The subplots A & B correspond to the two alternative methods of obtaining a NC graph with a single NC per phenotype: (A) Connecting all NCs of a given phenotype to a single node. (B) Keeping only the largest NC for each phenotype. We find that high-evolvability NC graphs remain high-evolvability NC graphs, even when the arithmetic mean or median is chosen.

##### S2.1.3 NC graph network analysis

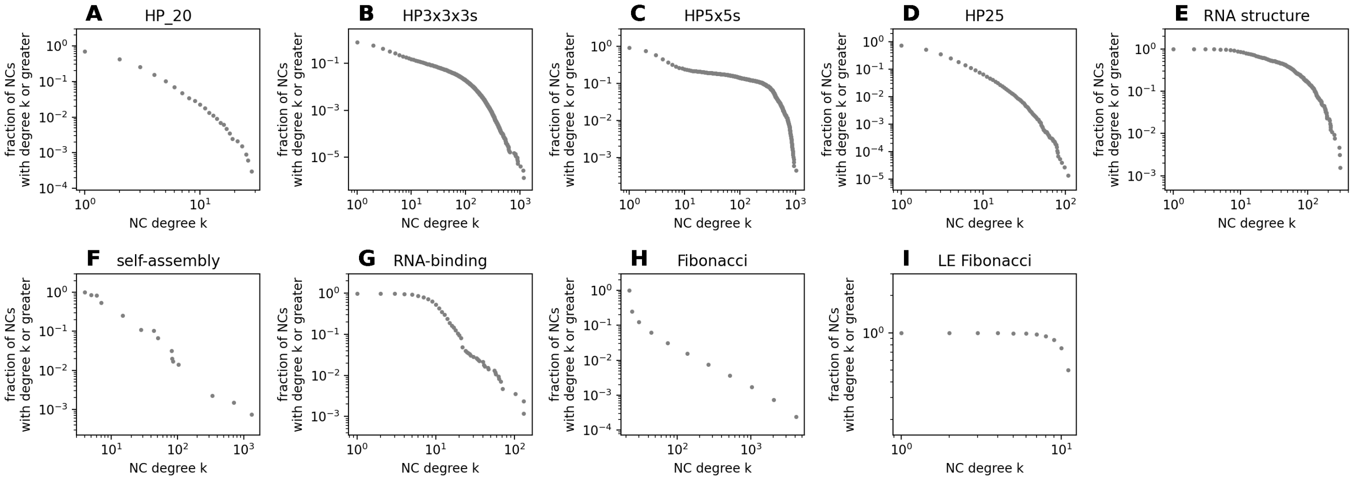

Figure S7: **Degree distribution of NC graphs:** For each NC graph, we plot the cumulative degree distribution, i.e. the fraction of nodes with degree $k$ or greater as a function of the degree $k$. In all networks, except the low-evolvability Fibonacci model, the degrees span several orders of magnitude.

Since NC graphs are at the centre of our analysis, we next turn to their network properties. First, we focus on the degree distribution of our NC graphs (Fig. S7). The network degree of a NC in its NC graph is closely related to its NC evolvability, but the two are not identical: for example, a NC of interest can be connected to two NCs of phenotype $p_x$, and this would contribute two units to its degree in the NC graph, but only one unit to its evolvability. We find that the NC graph degrees vary over several orders of magnitude, with a small number of high-degree nodes and many low-degree nodes (Fig. S7). This is not surprising given that evolvability and thus degree is correlated with NC size (see Figs. S3-S4). Some of these networks are approximately 'scale-free', i.e. their cumulative degree distributions in Fig. S7 follow a power law (see ref [37] for a more detailed definition).

Having quantified *how many* NCs each NC is connected to, let us focus on *which* NCs it tends to be connected to. In particular, we analyse whether high-degree nodes are preferentially connected to other high-degree nodes (Fig. S8), i.e. whether the network has degree assortativity. We find mixed results: our NC graphs have negative or small positive degree assortativity (again with the LE Fibonacci model as an outlier). Our finding does not invalidate previous observations of a positive degree assortativity in GP maps [38] since these previous results focused on a different scale: the network of genotypes in a NC, rather than the networks of NCs in the full GP map.

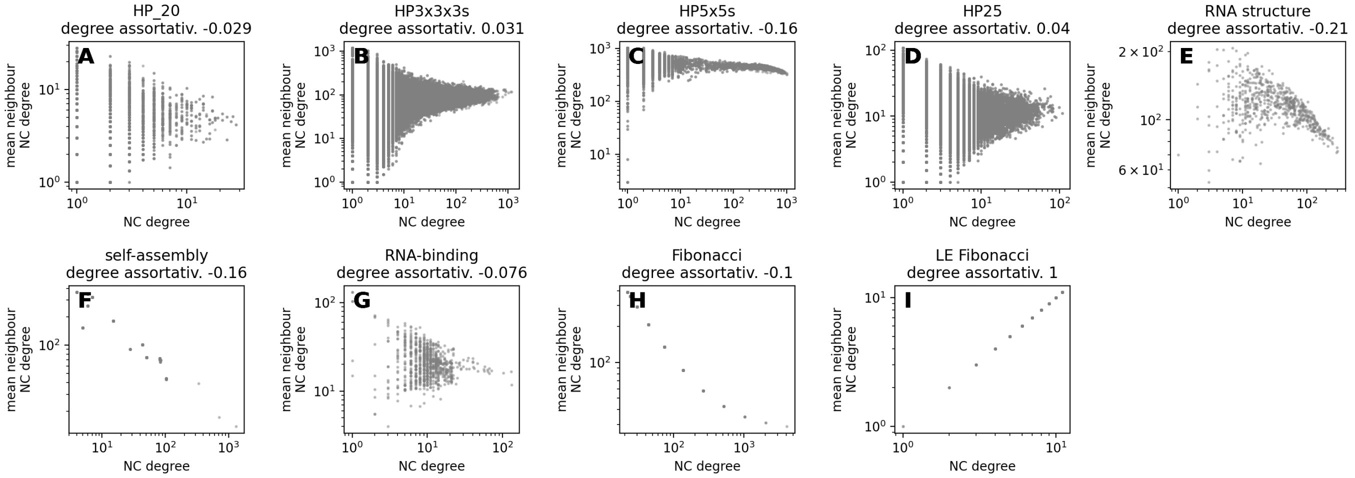

Figure S8: **Degree assortativity in NC graphs:** we take each node in a NC graph, and plot the mean degree of its neighbouring nodes against its own degree. We summarise this data by computing the degree assortativity coefficient with NetworkX [36] (see plot titles). NC graphs tend to have low or negative assortativity (again the low-evolvability Fibonacci model is an outlier by definition).

#### S2.2 Additional ruggedness analyses

In this section, we will provide additional data related to the ruggedness analysis in Fig. 2 of the main text.

##### S2.2.1 Changes to the underlying GP maps, including dimensionality and correlations

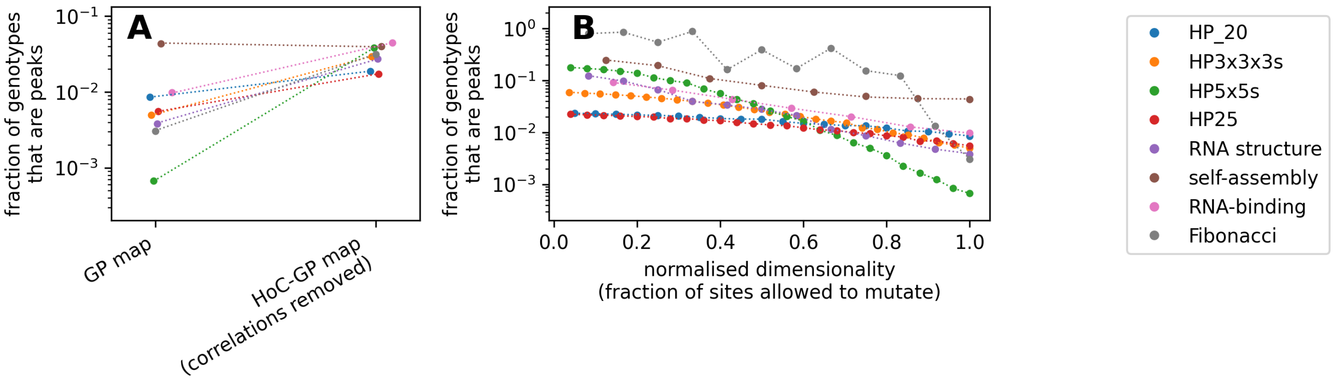

Figure S9: **Perturbing the correlations and dimensionality in a GP map indirectly affects ruggedness by altering the numbers, sizes and evolvabilities of NCs:** (A)We compare the ruggedness in each GP map to that of a corresponding zero-correlation GP-HoC map (the 'random GP map' in [14]). (B) We compute the ruggedness in reduced-dimensionality GP maps, where only a fraction of all $L$ sites are allowed to mutate (see ref [6]). In both plots, scatter points denote the mean ruggedness of each map in simulations. Dotted lines illustrate the prediction based on NC sizes and evolvabilities (eq. (2)), which indicate that changes in the set of NC sizes and evolvabilities fully account for changes in ruggedness. Each data point is based on a single GP map (for example, a single HoC-GP map, or a reduced-dimensionality GP map with a fixed set of mutating sites). Computational ruggedness estimates are based on $10^3$ PF maps for each GP map. Slight x-axis offsets are used in (A) to show overlapping points more clearly. The LE Fibonacci model is not shown because its construction already relies on excluding certain mutations.

One key takeaway from the calculations in section S1.1 is that the mean ruggedness depends only on the set of all NC sizes $|NC|$ and evolvabilities $\epsilon_{NC}$. Other GP map features only play an indirect role through their correlation with these properties. In this section, we test this indirect dependency by revisiting an analysis from Greenbury et al. [6] and varying two GP map characteristics: (1) The *dimensionality*, i.e. the number of positions allowed to mutate per genotype, and (2) *genetic correlations*, i.e. the clustering of genotypes of the same phenotype [14]. This clustering means that two mutationally connected genotypes are more likely to map to the same phenotype than in the corresponding uncorrelated null model. Thus, genetic correlations are closely linked to the high prevalence of neutral mutations in GP maps, and the formation of large NCs [14].

To investigate the role of these two GP map features, we follow Greenbury et al. [6, 14] and perturb them in two ways: (1) The dimensionality can simply be reduced by limiting the number of mutating sites in a genotype [6]. (2) The genetic correlations can be removed by building an uncorrelated "GP-HoC model" for a given GP map [14], by shuffling the assignment of phenotypes over genotypes. This was originally introduced as the “random GP map”, but here we use the term “GP-HoC model” to highlight parallels with uncorrelated House-of-Card models.

When we reduce the dimensionality and genetic correlations of our GP maps, we find that their ruggedness tends to increase (Fig. S9), as expected. These results, as well as the fact that the self-assembly model forms an exception due to its low number of phenotypes (see section S2.1.1), are in agreement with Greenbury et al. [6].

Having shown the computational ruggedness results, we turn to our ruggedness predictions (dotted lines in Fig. S9), which are based only on the set of NC sizes $|NC|$ and NC evolvabilities $\epsilon_{NC}$. These predictions continue to be accurate, indicating that the changes in NC sizes and evolvabilities fully predict the changes in mean ruggedness. These changes in NC sizes $|NC|$ and NC evolvabilities $\epsilon_{NC}$ can arise from GP map perturbations in many ways: For example, reducing dimensionality implies reducing the number of allowed mutations, which can lower NC evolvabilities by removing mutations connecting two NCs. Alternatively, reducing the number of allowed mutations can fragment an initially connected NC into several lower-evolvability NCs. Similarly, the reduction of genetic correlations through randomisation can break NCs into several lower-evolvability NCs. Thus, concepts like dimensionality and genetic correlations are indeed important for the ruggedness in GPF maps, but this effect is an indirect one mediated by their effects on the numbers, sizes and evolvabilities of NCs.

##### S2.2.2 The impact of deleterious phenotypes and zero-evolvability NCs on ruggedness

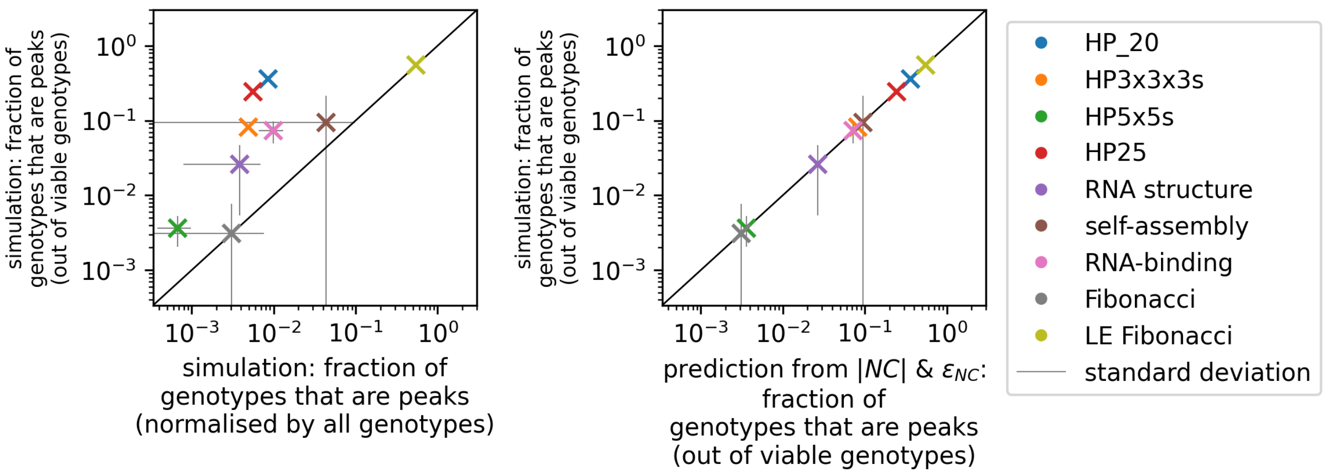

Figure S10: **Prevalence of peaks normalised relative to the set of all ‘viable’ genotypes (i.e. not mapping to the deleterious phenotype):** (A) Simulation data comparing the renormalised fraction of peaks (y-axis) to the fraction relative to all genotypes (x-axis, this is the data from the main text). (B) The renormalised fraction of genotypes in peaks (y-axis) is well-predicted by NC sizes $|NC|$ and NC evolvabilities $\epsilon_{NC}$ using eq. (18) (x-axis).

Finally, we analyse the impact of two GP map peculiarities on ruggedness estimates: the presence of ‘deleterious’ phenotypes and the existence of zero-evolvability NCs.

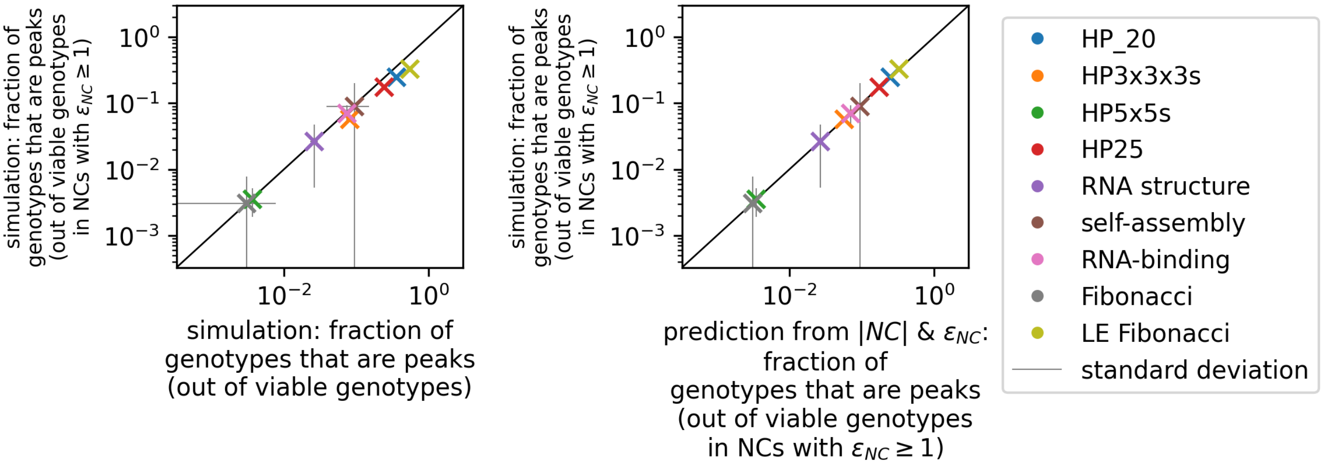

Figure S11: **Prevalence of peaks relative to the number of genotypes in viable & non-isolated NCs (with $\epsilon_{NC} > 0$):** (A) Simulation data comparing the renormalised fraction of peaks among viable, non-isolated NCs (y-axis) to the fraction relative to all viable NCs (x-axis, this is the data from Fig. S10). (B) The renormalised fraction of genotypes in peaks (y-axis) is well-predicted by NC sizes $|NC|$ and NC evolvabilities $\epsilon_{NC}$ using eq. (19) (x-axis).

Qualitatively distinct phenotypes exist in all of our GP maps: For example, non-uniquely folding sequences in the HP model, non-deterministic or non-finite outcomes in the self-assembly model and structures without base pairs in the RNA secondary structure [6] (however, for the case of RNA, they are thus much less prevalent for longer, more biologically relevant sequence lengths [20]). Similarly, some sequences in the RNA-binding model do not bind a single RNABP, and genotypes in the Fibonacci model are 'undefined' if they lack a stop codon [12]. Throughout this paper, we follow the worst-case treatment of Greenbury et al. [6], and treat all genotypes with these criteria as deleterious and of zero fitness. Thus, they cannot form accessible paths or escape routes from peaks, and are excluded from the NC graph.

Since genotypes mapping to the deleterious phenotype cannot form peaks, it may be useful to exclude them from the ruggedness normalisation and instead normalise the ruggedness of our GP maps relative to the number of 'viable' genotypes $\sum_{\text{NC indices } i} |NC_i|$ as follows (i.e. genotypes mapping to any phenotype except the deleterious phenotype):

$$\langle \text{ruggedness} \rangle = \frac{1}{\sum_{\text{NC indices } i} |NC_i|} \sum_{\text{NC indices } i} \frac{|NC_i|}{\epsilon_i + 1} \tag{18}$$

Fig. S10 shows that the renormalised mean ruggedness is well-predicted by eq. (18).

Next, we turn to zero-evolvability NCs, which only have the deleterious phenotype in their mutational neighbourhood. They technically form a peak under any PF map as long as the deleterious phenotype is treated as zero-fitness. However, such peaks may not be relevant for evolutionary processes, since such mutationally isolated NCs cannot be reached by mutations, and are unlikely to arise as initial conditions. We therefore further normalise our predicted and simulated number of peaks relative to all viable genotypes that are not part of zero-evolvability NCs (Fig S11):

$$\langle \text{ruggedness} \rangle = \left( \sum_{\text{NCs indices with } \epsilon_i > 0} |NC_i| \right)^{-1} \times \sum_{\text{NCs indices with } \epsilon_i > 0} \frac{|NC_i|}{\epsilon_i + 1} \tag{19}$$

In Fig. S11, we see that additionally excluding zero-evolvability NCs has a minor impact on our results because zero-evolvability NCs tend to be small with low $|NC_i|$ (see table. 1), with the exception of the low-evolvability Fibonacci model. Moreover, the renormalised mean ruggedness continues to be well-predicted by NC evolvabilities and sizes (eq. (19)).

We note that drawing the line between 'viable' and 'deleterious' may be more difficult in more complex biological GP maps. The point here is that our formalism can still take unviable mutations into account.

##### S2.3 Additional analyses on peak heights

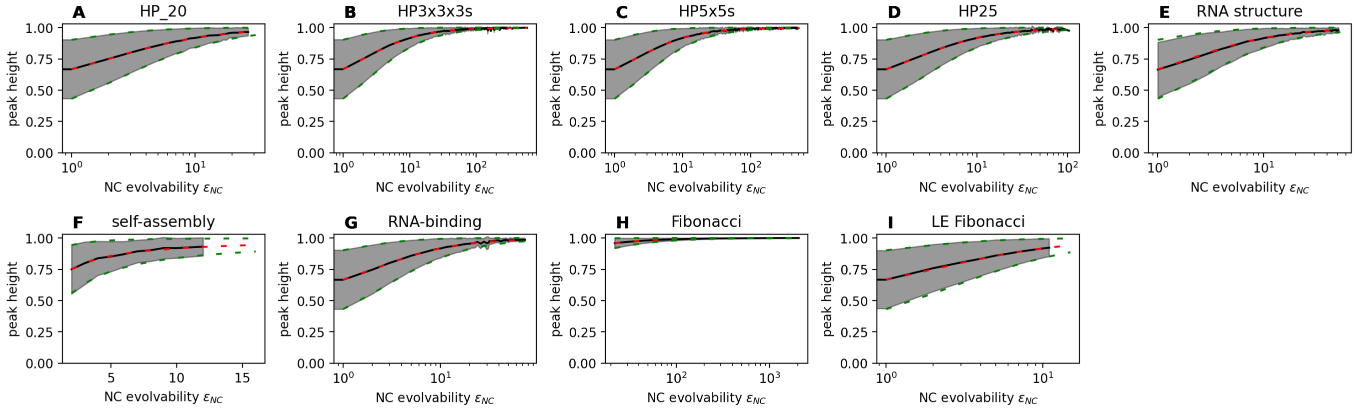

Figure S12: **More evolvable peaks have higher fitness:** each subplot shows data for one GP map. We simulate GPF maps sampled from a uniform distribution between 0 and 1, and plot peak fitness against evolvability (mean in grey and standard deviation as shaded grey area). In agreement with the analytic predictions from eqs (4 & 5) (red and green), high-evolvability peaks tend to be high in fitness. The simulation results are based on $10^3$ PF map realisations for each GP map, with up to $5 \times 10^6$ peaks per GP map included in the plot.

In section S1.2, we analytically derived the expected fitness of a peak NC as a function of its NC evolvability $\epsilon_{NC}$. We found that high-evolvability NCs tend to form high-fitness peaks, if they are peaks at all. This relationship was tested only for the RNA structure map in the main text, and so data for all GP maps is shown in Figs S12 & S13. We find that the analytic predictions are in excellent agreement with simulation data. Interestingly, for the exponential distribution (Fig. S13), mean fitness continues to increase with NC evolvability, without the saturation seen for the uniform distribution, approximately as $\int_1^{\epsilon_{NC}+1} (\lambda x)^{-1} dx = \log(\epsilon_{NC} + 1)/\lambda$. This is simply because there is no upper limit to the possible fitness values in the exponential distribution. This means that the higher the evolvability $\epsilon_{NC}$, the more neighbours need to be outcompeted, and thus the higher the expected fitness if the NC is a peak.

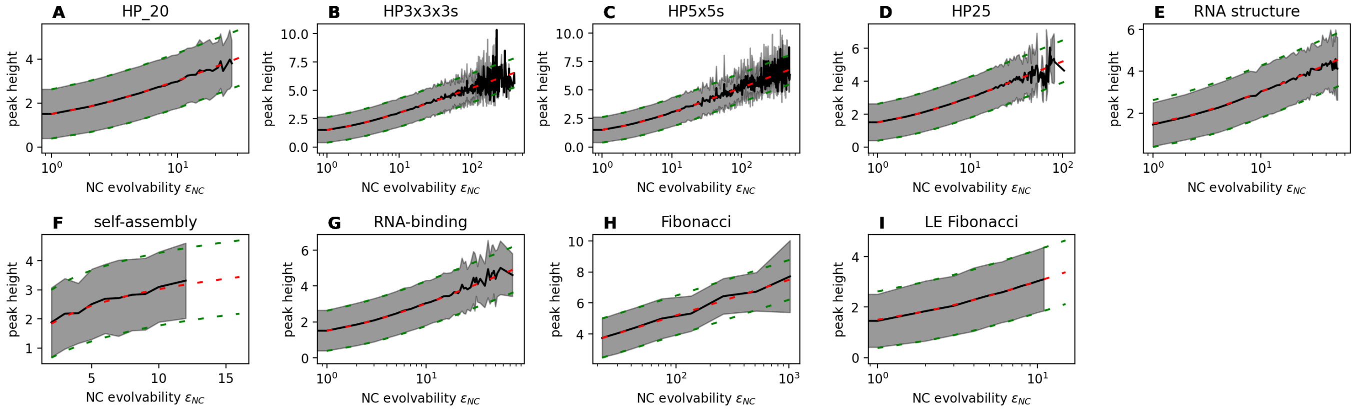

Figure S13: **More evolvable peaks are higher:** here we repeat the analysis from Fig. S12, but for a PF map sampled from an exponential distribution with rate parameter $\lambda = 1$. Again, we find that higher-evolvability peaks tend to be of higher fitness, in agreement with the prediction of eqs (6 & 8).

The relationship between peak evolvability and height also implies an indirect link between peak size $|NC|$ and height: larger NCs tend to have higher evolvability (see Fig S4) and thus tend to form higher peaks. This is confirmed in Figs S14 & S15. In all GP maps except the low-evolvability Fibonacci model, larger peaks tend to be of higher fitness. This trend is weakest in the self-assembly GP map since even large NCs in this map only reach moderate evolvabilities (see Fig S4), as well as in the RNA-binding map, where the range of evolvabilities and NC sizes is limited, probably partly due to the short sequence length of only $L = 7$.

The low-evolvability Fibonacci model is an exception precisely because the link between peak size and height is an indirect one: larger peaks are of higher fitness only if NC evolvability is *positively* correlated with NC size, which is not the case in the low-evolvability Fibonacci model (see Fig. S4). While the low-evolvability Fibonacci model is useful for elucidating this relationship and disentangling direct and indirect links, it is not a realistic GP map. Realistic GP maps are thought to have a positive relationship between size and evolvability [15, 33], implying that they will have a positive correlation between peak size and evolvability.

The intuitive link between neighbourhood sizes (here evolvabilities) and peak heights holds beyond GPF maps: for example, in the HoC model, peaks tend to be higher for landscapes with larger mutational neighbourhoods [39]. However, there is one key difference: In the HoC model, the neighbourhood size is simply set by the sequence length $L$ and alphabet size $K$, and thus takes a single value for an entire landscape. In GP maps, however, the relevant neighbourhood size is the evolvability, and this can differ by orders of magnitude *within* a single landscape, creating a rich phenomenology with peaks of different heights, evolvabilities and sizes.

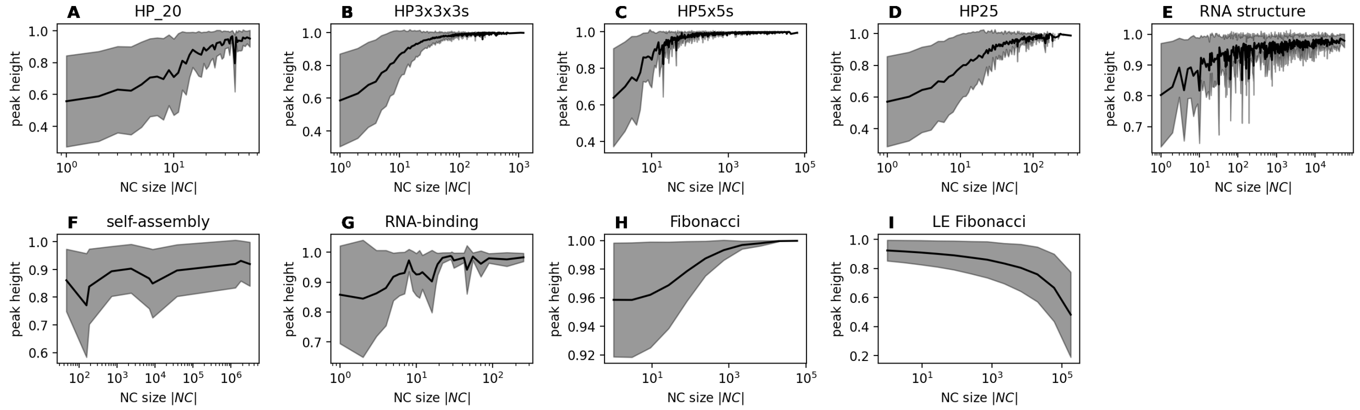

Figure S14: **Larger peaks tend to be higher due to their higher evolvability:** here we show the peak heights from Fig. S12 (for a PF map drawn from a uniform distribution between 0 and 1), but as a function of the NC size rather than NC evolvability. Larger NCs, if they are peaks, tend to be higher. This is an indirect effect: larger NCs tend to have higher evolvability (Fig. S4) and higher-evolvability peaks tend to be higher (eq. (4)). Because the positive correlation depends on the NC-size-evolvability relationships, the trend is negative for the low-evolvability Fibonacci model, where larger NCs are less evolvable (see Fig. S4).

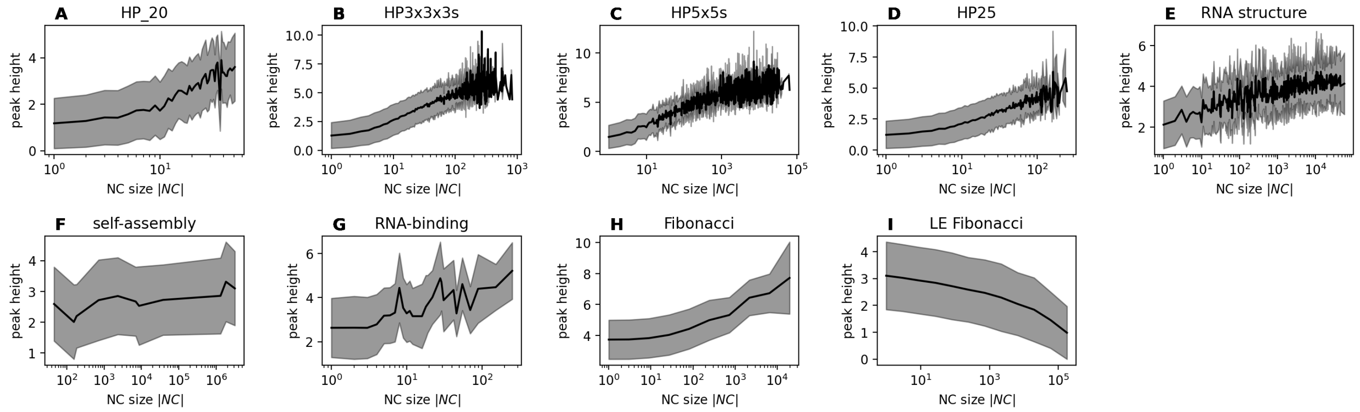

Figure S15: **Larger peaks tend to be higher due to their higher evolvability:** same as Fig. S14, but for a PF map drawn from an exponential distribution with rate parameter $\lambda = 1$.

#### S2.4 Additional navigability analyses

##### S2.4.1 Navigable vs. non-navigable synthetic NC graphs

In this section, we use synthetic networks to test our navigability scaling from eq. (11) on NC graphs with a variety of evolvability distributions, including broader distributions than assumed in the derivation. We will do this with the simplification of NC graphs with one NC per phenotype, where network degree is the same as evolvability.

To create a test set of NC graphs, we built synthetic networks with degrees drawn from three different types of distributions: a single degree value (i.e. a delta-function-like distribution), a Poisson distribution and a geometric distribution. For each type of distribution, we varied the number of draws and the mean, to design NC graphs with different numbers of nodes ('numbers of phenotypes' in this context) and mean degree ('mean evolvability' in this context). Specifically, we used the parameters (50, 100, 200, 300, 500, $10^3$) for the number of phenotypes and ($0.005n_p$, $0.015n_p$...$0.245n_p$) for the mean degree. Once we had created a suitable list of node degrees, and after checking that the degree sum is even to ensure that a solution exists, we fed this into the configuration_model() function of the NetworkX Python package to create a network with the desired degree distribution. While the resulting network satisfies the specified degree list, it may contain multi-edges and self-loops, which we removed. Finally, we discarded any isolated nodes to avoid artefacts.

Fig S16 gives an overview of the properties of the resulting networks and their navigabilities, with each row corresponding to one type of evolvability distribution. First, we analysed the evolvability distributions of the constructed networks (columns I & II in Fig S16, with one row for each type of distribution). Then, we investigated their navigabilities (column III in Fig S16). We find that low-evolvability NC graphs are non-navigable, as expected from eq. (11) (black line), even though the assumption of a single-peaked degree distribution around a well-defined mean $\epsilon_{NC}$ is not met for the NC graphs based on geometric distributions.

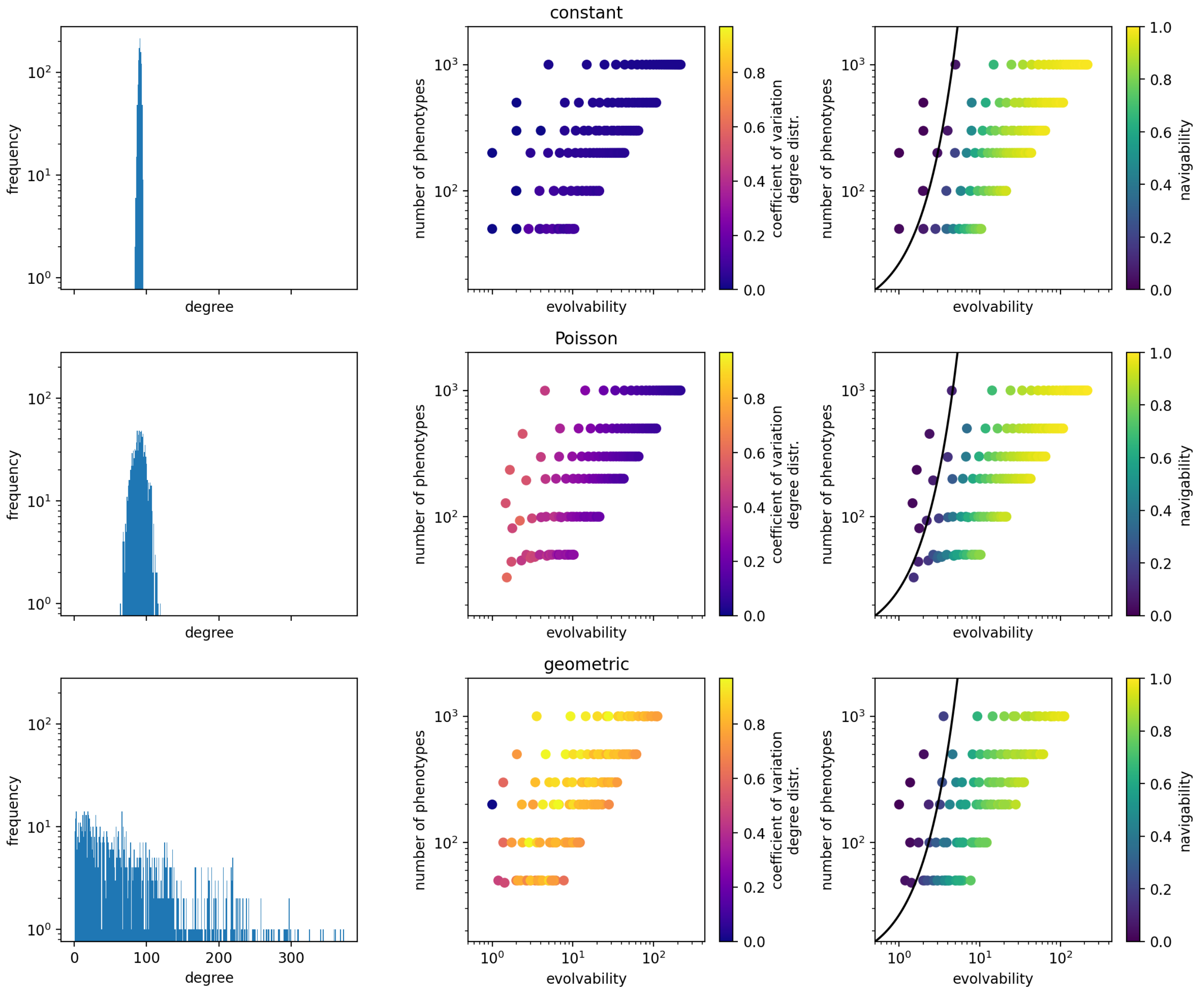

Figure S16: **Test of navigability scaling on synthetic networks:** Here we use a test set of synthetic NC graphs with degrees sampled from three different distributions: a single degree value, a Poisson distribution and a geometric distribution (one per row), with a variety of mean degree and number of nodes in each case. Columns I & II characterise these synthetic networks: column I shows the degree distribution for one example NC graph per distribution type. Column II plots the number of phenotypes (nodes) in each NC graph against its evolvability (geometric mean of the degree distribution), illustrating that a diverse range of networks was created for each distribution. The colour of the scatter point is given by the coefficient of variation of the NC graph's degree distribution, illustrating the difference between the different types of distributions. Finally, in the third column, the navigability of each network is indicated by the colour (based on $10^2$ PF maps and 10 source-target pairs each). We find that low-evolvability NC graphs are non-navigable, as expected from eq. (11) (black line).

##### S2.4.2 Navigable vs. non-navigable GP maps in a sample of GP maps based on the Fibonacci model

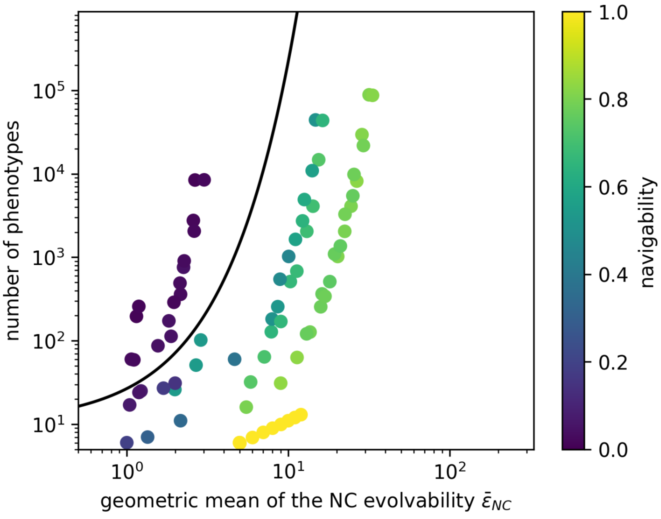

Figure S17: **Navigable vs. non-navigable GP maps in a sample of GP maps based on the Fibonacci model:** For each Fibonacci-based GP map (see text), we plot the number of phenotypes against the geometric mean of its NC evolvabilities, and indicate its navigability by the colour scale (based on $10^2$ realisations of the PF map with 10 source-target pairs each). We find that low-evolvability GP maps are non-navigable, as expected from eq. (11) (black line), despite the approximations made in the derivation.

As a second test of how well our scaling from eq. (11) identifies non-navigable GP maps, we apply it to an additional test set of GP maps. Ideally, these GP maps fulfil our assumption of having a single NC per phenotype but differ in their numbers of phenotypes and evolvabilities. Thus, we work with variations of the Fibonacci model, which is simple and has a single NC per phenotype. To generate GP maps with different numbers of phenotypes, we vary their sequence length and alphabet size ($5 \leq L \leq 13$ and $2 \leq K \leq 5$, except maps with L/K combinations of 12/4, 13/4, 10/5, 11/5, 12/5, 13/5, which were computationally unfeasible). To create even more maps, we replaced increasing fractions of each GP map with 'deleterious' genotype: We created four derivative maps from each GP map, keeping a set fraction of all phenotypes in each case ($10^{-3}$, $10^{-2}$, $10^{-1}$ and 0.5) and replacing the genotypes corresponding to the remaining phenotypes with 'deleterious/unviable' genotypes. Thus, the removed phenotypes disappear from the NC graph and all derived metrics (number of phenotypes $n_p$ and the NC evolvability distribution). If any NC has zero evolvability $\epsilon_{NC} = 0$ after this procedure, we also delete this phenotype since zero-evolvability NCs could affect navigability in non-trivial ways. We only analyse GP maps with more than five nodes in their NC graph to avoid small-size effects.

Thus, we created GP maps with different evolvabilities and number of phenotypes, and computed their navigabilities through simulations. Fig S17 shows that this test set includes both navigable and non-navigable GP maps with a range of evolvabilities and numbers of phenotypes. Among these GP maps, our approximate scaling from eq. (11) successfully identifies a minimum evolvability, below which GP maps are non-navigable.

##### S2.4.3 Disjoint components and navigability in NC graphs

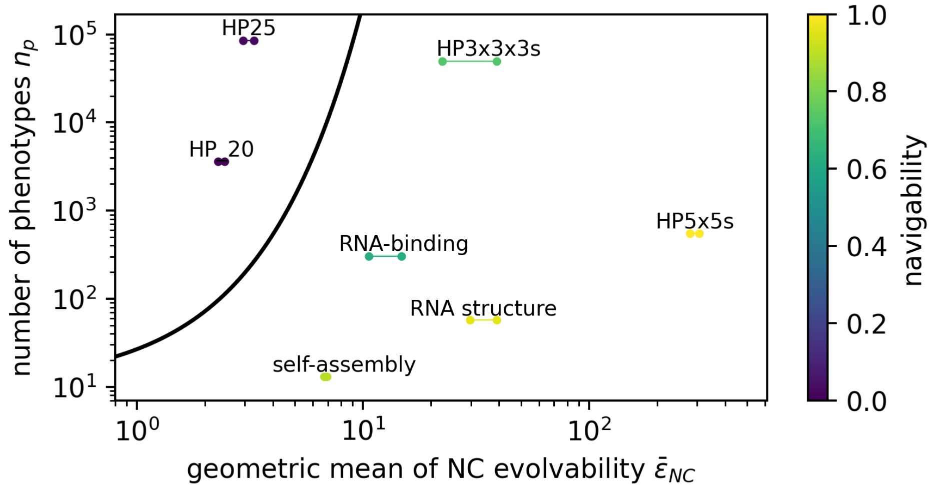

Figure S18: **Evolvability plays a key role for navigability, even when working only with the largest connected component of a NC graph:** First, every NC graph is reduced to its largest connected component, to exclude source/target pairs, where no mutational path exists (neither accessible nor non-accessible paths). Navigability is computed on this connected NC graph. As in Fig. 4 the main text, the evolvability of each GP map is represented by two data points, corresponding to two ways of resolving NC fragmentation. For each GP map, the number of phenotypes is plotted against the geometric mean of the evolvabilities, with navigability shown as a colour. As before, eq. (11) (black line) successfully identifies an evolvability below which GP maps are non-navigable, despite relying on strong approximations.

Finally, we will investigate one potential confounding factor in the navigability analysis in the main text: the presence of disconnected, zero-evolvability NCs, which exist in several GP maps (see table 1), and cannot have accessible paths to any target phenotypes. To exclude these NCs are possible confounding factors, we reduce each NC graph to its largest connected component (largest in terms of the highest number of NCs). In this setting, any NC has a mutational path to any other NC, but this path does not have to be an accessible, fitness-increasing path (see, for example [40] for low-navigability fitness landscapes even without fragmentation into disconnected components) .

Even after reducing each NC graph to a single connected component, the navigabilities and evolvabilities of our GPF maps differ drastically (Fig. S18): in two of the HP protein models (HP_20 and HP25), evolvabilities remain low, indicating that each NC is only connected to a small number of different phenotypes. As expected from our approximate scaling (eq. (11)), these low-evolvability GPF maps are of low navigability even when they do not have mutationally disjoint components. The remaining GPF maps have higher evolvabilities and higher navigabilities, as before.
